# A fully joint Bayesian quantitative trait locus mapping of human protein abundance in plasma

**DOI:** 10.1101/524405

**Authors:** Hélène Ruffieux, Jérôme Carayol, Radu Popescu, Mary-Ellen Harper, Robert Dent, Wim H. M. Saris, Arne Astrup, Jörg Hager, Anthony C. Davison, Armand Valsesia

## Abstract

Molecular quantitative trait locus (QTL) analyses are increasingly popular to explore the genetic architecture of complex traits, but existing studies do not leverage shared regulatory patterns and suffer from a large multiplicity burden, which hampers the detection of weak signals such as *trans* associations. Here, we present a fully multivariate proteomic QTL (pQTL) analysis performed with our recently proposed Bayesian method LOCUS on data from two clinical cohorts, with plasma protein levels quantified by mass-spectrometry and aptamer-based assays. Our two-stage study identifies 136 pQTL associations in the first cohort, of which > 80% replicate in the second independent cohort and have significant enrichment with functional genomic elements and disease risk loci. Moreover, 78% of the pQTLs whose protein abundance was quantified by both proteomic techniques are confirmed across assays. Our thorough comparisons with standard univariate QTL mapping on (1) these data and (2) synthetic data emulating the real data show how LOCUS borrows strength across correlated protein levels and markers on a genome-wide scale to effectively increase statistical power. Notably, 15% of the pQTLs uncovered by LOCUS would be missed by the univariate approach, including several *trans* and pleiotropic hits with successful independent validation. Finally, the analysis of extensive clinical data from the two cohorts indicates that the genetically-driven proteins identified by LOCUS are enriched in associations with low-grade inflammation, insulin resistance and dyslipidemia and might therefore act as endophenotypes for metabolic diseases. While considerations on the clinical role of the pQTLs are beyond the scope of our work, these findings generate useful hypotheses to be explored in future research; all results are accessible online from our searchable database. Thanks to its efficient variational Bayes implementation, LOCUS can analyse jointly thousands of traits and millions of markers. Its applicability goes beyond pQTL studies, opening new perspectives for large-scale genome-wide association and QTL analyses.

**Author summary:** Exploring the functional mechanisms between the genotype and disease endpoints in view of identifying innovative therapeutic targets has prompted molecular quantitative trait locus studies, which assess how genetic variants (single nucleotide polymorphisms, SNPs) affect intermediate gene (eQTL), protein (pQTL) or metabolite (mQTL) levels. However, conventional univariate screening approaches do not account for local dependencies and association structures shared by multiple molecular levels and markers. Conversely, the current joint modelling approaches are restricted to small datasets by computational constraints. We illustrate and exploit the advantages of our recently introduced Bayesian framework LOCUS in a fully multivariate pQTL study, with ≈ 300K tag SNPs (capturing information from 4M markers) and 100 – 1,000 plasma protein levels measured by two distinct technologies. LOCUS identifies novel pQTLs that replicate in an independent cohort, confirms signals documented in studies 2 – 18 times larger, and detects more pQTLs than a conventional two-stage univariate analysis of our datasets. Moreover, some of these pQTLs might be of biomedical relevance and would therefore deserve dedicated investigation. Our extensive numerical experiments on these data and on simulated data demonstrate that the increased statistical power of LOCUS over standard approaches is largely attributable to its ability to exploit shared information across outcomes while efficiently accounting for the genetic correlation structures at a genome-wide level.

## Introduction

Questioning the genetic contribution to human diseases has become a critical step towards predicting health risks and developing effective therapies [1–3]. However the functional network of interacting pathways between the genotype and disease endpoints largely remains a “black box”, so the expected transformation of medicine has only begun. The analysis of endophenotypes such as gene, protein or metabolite levels, via molecular quantitative trait locus (QTL) studies may provide deeper insight into the biology underlying clinical traits [3]. While eQTL studies are now routinely performed, pQTL studies have emerged only recently [4–9]. These studies allow the exploration of the genetic bases of several diseases, as certain proteins may act as proxies for specific clinical endpoints [10]. However two major hurdles hamper pQTL analyses. First, owing to the number of tests that they entail, conventional univariate approaches lack statistical power for uncovering weak associations, such as *trans* and pleiotropic effects [11–13], while better-suited multivariate methods fail to scale to the dimensions of QTL studies. Second, the clinical data complementing QTL data are often very limited, restricting subsequent investigation to external information from unrelated populations, health status or study designs, and rendering some degree of speculation unavoidable.

In this paper, we demonstrate that both concerns can be addressed using statistical approaches and data tailored to the problem under consideration: we present an integrative genome-wide pQTL study of two clinical cohorts performed with our Bayesian joint QTL method LOCUS [14], which simultaneously accounts for all the genetic variants and proteomic outcomes, thereby leveraging the similarity across proteins controlled by shared functional mechanisms. We employ a two-stage design, using data from the Ottawa cohort (*n* = 1, 644) [15] for discovery and replicating our findings in the independent DiOGenes cohort (*n* = 789) [16]. Each dataset involves protein plasma levels quantified by mass-spectrometry (MS) and aptamer-based (SomaLogic, [17]) assays, which permits thorough cross- and intra-platform validation.

Our work aims to illustrate the utility and feasibility of multivariate Bayesian QTL analyses. Notably, we show the gain in statistical power achieved by LOCUS on our data by confronting the validated hits with those of a classical two-stage univariate design. We also assess the performance of both approaches in simulations mimicking real data conditions.

Pertinent interpretation of pQTL loci in the context of complex diseases hinges on a careful examination of metabolic and clinical parameters from the same subjects or, at a minimum, from a population presenting similar clinical characteristics. Using comprehensive data from the two pQTL cohorts, we evaluate the enrichment of the proteins under genetic control for associations with clinical parameters; this assessment suggests that our pQTL findings may ease the identification of proteomic biomarkers and calls for further dedicated research. All results are available from our online browser https://locus-pqtl.epfl.ch/db.

## Results

### Two-stage pQTL analyses

We analyzed the pQTL data from the Ottawa and the DiOGenes cohorts in a two-stage study. The DiOGenes cohort recruited overweight/obese, non-diabetic subjects, while the Ottawa study was led in a specialized obesity practice where subjects had severe obesity, dyslipidemia and insulin resistance disorders (S1 Table).

We used LOCUS ([14], Material and methods and Fig 1a–c) for joint analyses of both proteomic datasets from the Ottawa cohort, quantified by MS and the multiplexed aptamer-based technology SomaLogic [17], respectively. We adjusted all analyses for body mass index (BMI), age and gender. At FDR 5%, LOCUS identified 18 pQTL associations from the MS analysis, corresponding to 14 unique proteins and 18 SNPs, and 118 pQTLs from the SomaLogic analysis, corresponding to 99 proteins and 111 SNPs (S2 Table). We then undertook to replicate all uncovered pQTLs in the independent DiOGenes cohort, using MS and SomaLogic data. We validated 15 of the 18 discovered MS pQTLs, and 98 of the 118 discovered SomaLogic pQTLs at FDR 5% (S3 Table), yielding a replication rate of 83% in both cases (Fig 1d).

**Figure 1:**
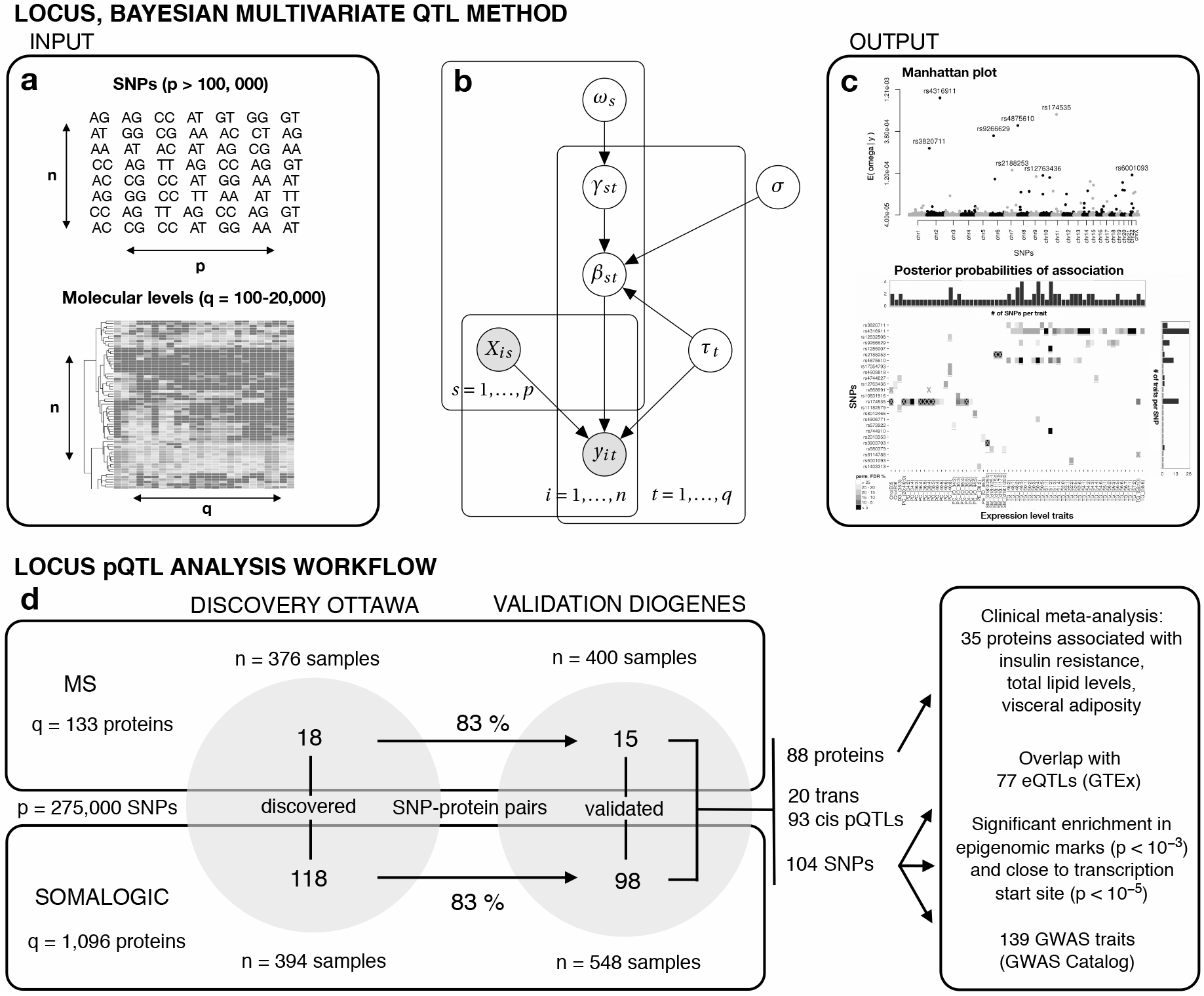
LOCUS model overview and study workflow. (a) Inputs to LOCUS are an *n × p* design matrix ***X*** of *p* SNPs, and an *n* × *q* outcome matrix ***y*** of *q* molecular traits, e.g., gene, protein, lipid, metabolite or methylation levels, for n individuals. The model accounts for all the SNPs and molecular traits jointly. (b) Graphical model representation of LOCUS. The effect between a SNP *s* and a traid *t* is modelled by *β_st_*, and γ_*st*_ is a latent variable taking value unity if they are associated, and zero otherwise. The parameqer *ω_s_* controls the pleiotropic level of each SNP, i.e., the number of traits with which it is associated. The parameter *σ* represents the typical size of effects, and the parameter *τ_t_* is a precision parameter that relates to the residual variability of each trait *t*. (c) Outputs of LOCUS are posterior probabilities of associations, pr(γ_st_ = 1 | ***y***), for each SNP and each trait (*p × q* panel), and posterior means for the pleiotropy propensity of each SNP, E (*ω_s_* | ***y***) (Manhattan plot). (d) Workflow of the pQTL study. The MS and SomaLogic pQTL data are analyzed in parallel. LOCUS is applied on the Ottawa data for discovery, and 83% of the 18 and 118 pQTL associations discovered with the MS and SomaLogic dath replicate in the independent study DiOGenes. The possible relevance ol hhe validated pQTLs for disease endpoints is explored via analyses of clinical parameters arom the Ottawa and DiOGenes cohorts. Further support is obtained by evaluating the overlap with eQTLs, epigenomic marks and GWAS risk loci.

We evaluated replication rates separately for *cis* and *trans* effects. With the MS data, all 15 *cis* Ottawa pQTLs replicated in DiOGenes, while the 3 *trans* pQTLs did not. With the SomaLogic data, 78 of 81 cis and 20 of 37 *trans* pQTLs could be validated (Fig 2c). The overall replication rates reached 97% for the cis pQTLs and 50% for the *trans* pQTLs; the *trans*-pQTL rate is in line with other pQTL studies [4, 7, 8]. Finally, 35 of our validated pQTLs are, to our knowledge, new, i.e., they do not overlap with pQTLs previously identified in the general population [4, 6–9], and this number drops to 20 using proxy search *r*^2^ > 0.8; four of these 20 hits have isoforms involved in known pQTLs, see S4 Table for the complete list.

**Figure 2:**
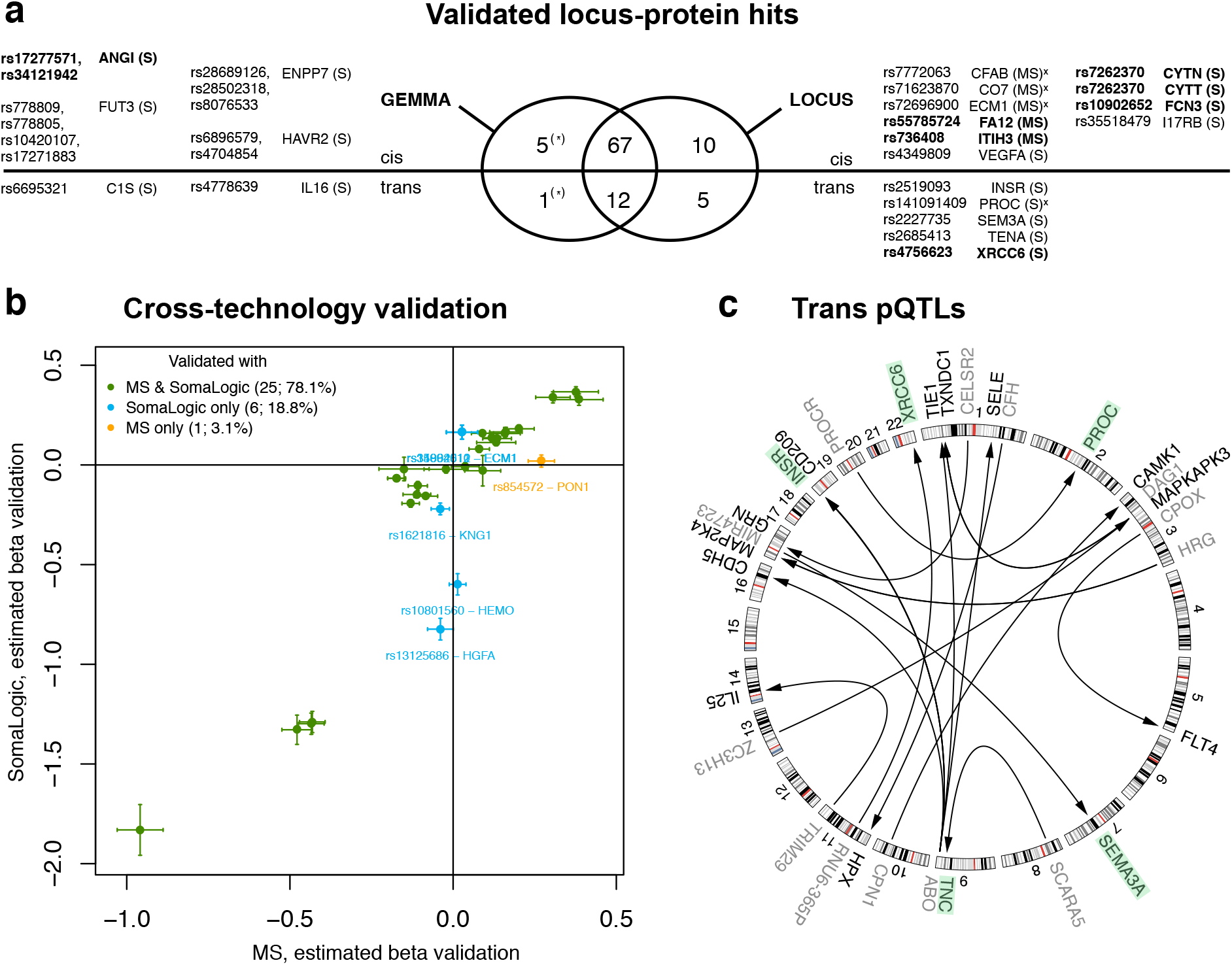
Overview of LOCUS validated pQTL hits. (a) Venn diagram for the locus-protein hits identified by the GEMMA and LOCUS two-stage analyses. The hits uncovered by GEMMA but not by LOCUS (left) and the hits uncovered by LOCUS but not by GEMMA (right) are listed; the stars indicate that the former were not tagged so not detectable by the LOCUS analyses. When multiple SNPs correspond to the same locus-protein hit, the SNP(s) with the top association(s) in the Ottawa cohort is/are shown. The novel hits (*r*^2^ > 0.8-proxy search) are in bold and the hits with dual replication in the alternative proteomic platform are marked with a cross (4 over the 4 quantified with both platforms). (b) Estimated effects (regression coefficients with standard errors) in DiOGenes for the validated pQTLs whose controlled protein is quantified by both technology (S3 Table). (c) Circular plot for the *trans*-pQTL associations uncovered by LOCUS (FDR < 5% for discovery and validation). Each arrow starts from the pQTL SNP with label indicating its closest gene (grey) and points to the gene (black) coding for the controlled protein. The proteins whose *trans* pQTLs are missed by GEMMA are highlighted in green.

### Cross-assay validation

The relative merits of MS untargeted and multiplexed aptamer-based techniques for quantifying protein abundance have been a subject of active debate over the past years [see, e.g., 18, 19]. On one hand, the former usually demonstrate very high specificity, but are limited in dynamic range and in their sensitivity to detect low abundance proteins. On the other hand, the latter – and in particular the recent SOMAscan technology provided by SomaLogic [17] – can target proteins across a wide dynamic range and include lowly expressed proteins not accessible to MS approaches, but are more subject to binding artefacts caused by the multiplexing. Hence, although rapidly improving, the two techniques remain complementary, and the combination of measurements by both approaches offers a convenient balance between specificity and sensitivity.

In the Ottawa and DiOGenes studies, a subset of 72 proteins was quantified by both the MS and SomaLogic techniques (S1 Appendix). For the pQTLs whose protein had dual measurement, this enabled an additional layer of validation with the alternative assay, on top of direct within-assay validation. Eight of the MS pQTLs could be assessed with SomaLogic (i.e., had protein levels available), and 7 of them replicated at FDR 5%. Likewise, of the 20 SomaLogic associations having MS measurements, 14 were confirmed (S3 Table). Moreover, the cross-assay correlation of estimated effects for the pQTLs with successful dual validation was high, *ρ* = 0.93, providing further support to our findings (Fig 2b).

Specificity issues and experimental platform-inherent variability might explain why the remaining seven pQTLs could not be confirmed using the alternative technology. Indeed, although most protein expression patterns quantified by the two technologies were in good agreement, some were not, in particular, the between-technology correlation for the genetically-driven protein levels HEMO, HGFA, LYSC and PON1 was lower than 0.25 in the two cohorts (S1 Appendix). In line with this hypothesis, recent work by Kim and colleagues [19] concluded on an overall good reproducibility with SomaLogic, yet with some non-negligible technical variability (intra-class correlation coefficient ≈ 0.4 for 91 of the proteins that they tested).

### Comparison with a standard univariate pQTL analysis

There is a broad consensus in the biostatistical community about the benefits of collectively accounting for the SNPs while leveraging information across multiple correlated outcomes [e.g., 20–22]. However, many also acknowledge the mathematical and computational difficulties hampering these practices in genome-wide association or QTL studies [e.g., 23, 24]. Pilot simulation experiments using LOCUS [14] have indicated that its flexible hierarchical sparse regression model coupled with a scalable variational inference scheme (Fig 1a–c) can effectively increase statistical power at the scale required by current molecular QTL studies. Here we provide sound practical evidence – in the context of the Ottawa and DiOGenes studies and based on thorough comparisons with the classical univariate screening design – that this tailored multivariate framework can lead to higher replication rates and novel pQTL discoveries. Namely, we compare LOCUS with the univariate linear mixed model approach GEMMA [25, 26] in two complementary ways. First, we use synthetic data to repeatedly evaluate their variable selection performance for a series of data scenarios where the ground truth (here QTL associations) is known since simulated. Second, we confront the findings (number of *cis/trans* locus-protein hits detected, replication rates) of LOCUS two-stage analysis to those obtained by re-analyzing the Ottawa and DiOGenes datasets with GEMMA.

### Simulations

Simulation studies can be far from reflecting real data scenarios. To best emulate our study and obtain a realistic assessment of the relative performance of the multivariate and univariate approaches, special care was taken to use synthetic data that resemble the real data and embody accepted principles of population genetics (Material and methods). In particular, we employed real SNPs from our study as candidate predictors and we simulated proteomic levels by replicating the block-correlation structure and variability of the 133 MS protein levels, using our freely available data generator `echoseq`. A comparison of the synthetic and real proteomic levels is provided in S2 Appendix.

The first simulation study uses the SNPs from chromosome one of the Ottawa cohort and the 133 outcomes mimicking the MS proteomic levels, enforcing that the SNPs explain together at most 35% of each protein variance (Material and methods). The ROC curves (Fig 3a) show a net gain in power for selections with LOCUS compared to GEMMA. The average standardized partial areas under the curve (pAUC) with 95% confidence intervals are 0.926 ± 0.005 for LOCUS and 0.840 ± 0.005 for GEMMA. The second simulation study generalizes this observation to a grid of data generation scenarios with different effect sizes and numbers of proteins under genetic control: the average standardized pAUCs (Fig 3b) are greater for LOCUS than for GEMMA in most cases and are comparable for very low proportions of protein variance explained by the SNPs.

**Figure 3:**
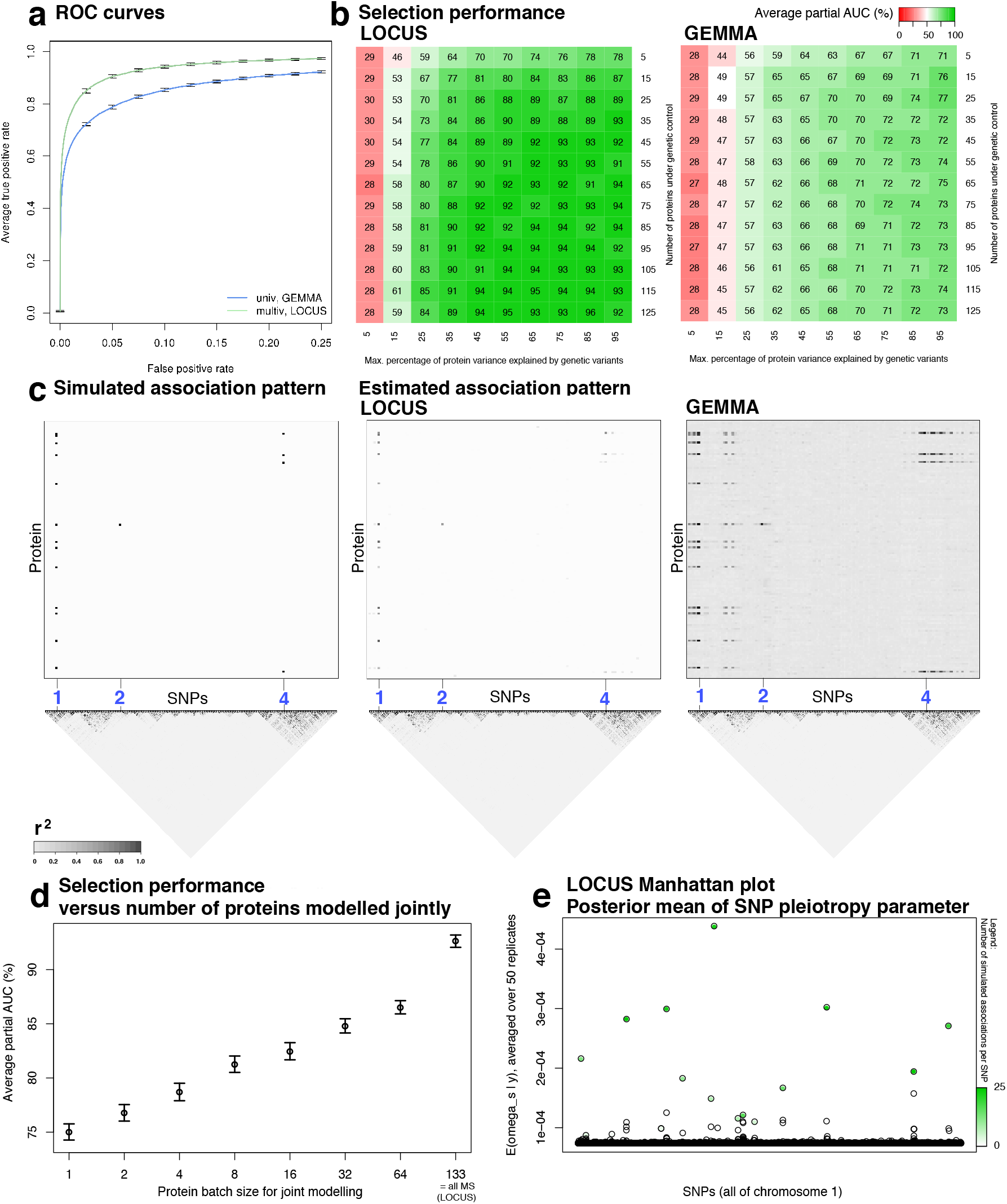
Selection performance of LOCUS and GEMMA on simulated data. (a) Truncated average ROC curves with 95% confidence intervals (50 replicates) for the detection of SNP-trait associations. (b) Average standardized partial AUC of LOCUS and GEMMA for a grid of effect sizes (*x*-axis) and signal sparsity (*y*-axis, 20 replicates for each scenario). (c) Simulated association pattern and patterns recovered by LOCUS and GEMMA, averaged over the 50 replicates. The plots display a window of 350 SNPs (*x*-axis) containing the first three SNPs with simulated associations (blue labels), along with their LD pattern. (d) Average standardized AUC with 95% confidence intervals for different numbers of proteins modelled jointly, i.e., the 133 simulated proteins are randomly partitioned into batches of size *q*_0_ (*x*-axis) and LOCUS is applied separately on these batches: *q*_0_ = 1 corresponds to modelling the proteins one by one and *q*_0_ = 133 corresponds to modelling all 133 proteins jointly, as achieved by a classical application of LOCUS. (e) Posterior mean of the LOCUS parameter representing the propensity for SNPs to be pleiotropic. Its magnitude satisfactorily reflects the number of simulated associations per SNP (color-coded).

By design, univariate screening approaches do not exploit association patterns common to multiple outcomes or markers; they analyze the outcomes individually, and do not account for linkage disequilibrium (LD), which often results in a number of redundant proxy discoveries at loci with strong LD (Fig 3c). At a given FDR, this hampers the detection of weak but genuine signals because of the large multiple testing burden. Post-processing strategies accounting for local LD structures exist [27, 28] and alleviate the problem to some extent. Instead of applying such corrections, LOCUS anticipates and addresses the question by enforcing sparsity directly at the modelling stage. It better discriminates truly associated SNPs from their correlated neighbours (Fig 3c), owing to its simulated annealing procedure. We do not claim that LOCUS can provide direct information on the regulatory potential of individual variants in real data settings, as this always requires follow-up studies at the level of loci, e.g., using fine-mapping analyses or in-lab experiments. We do argue however that its effective handling the LD structures can improve the selection of candidate variants for these subsequent functional studies, which may save substantial research investment.

Finally, the improvement over GEMMA and other classical approaches also comes in large part from the capacity of LOCUS to handle all correlated outcomes jointly, thereby borrowing strength across them. This is illustrated in our third simulation study (Fig 3d), whereby LOCUS is applied separately to multiple outcome batches of increasing sizes, from one protein per batch, corresponding to a univariate treatment of each proteomic outcome, to all 133 simulated proteins, corresponding to a regular application of our method. The AUCs indicate a significant increase of power when the outcomes are modelled jointly, as shared information across them can be effectively exploited to enhance the detection of weak effects. Technically, this shared information feature is mainly achieved via a specific model parameter, *ω_s_* (Fig 1b), which directly reflects the propensity of each SNP to be pleiotropic, i.e., to be associated with multiple outcomes (Fig 3e); we will return to the practical value of this parameter in the discussion of a pleiotropic locus identified in the real data study. Finally, we emphasize that our fully joint analysis is only possible thanks to the efficient variational inference implementation of LOCUS which supersedes the prohibitively slow Markov chain Monte Carlo (MCMC) routines traditionally employed for Bayesian inference.

These simulations tailored to the real data at hand prefigure the gain of power achieved by LOCUS over standard methods for our pQTL study, as we next discuss.

### Performance on the real data

We now compare LOCUS and GEMMA on the real data under the typical two-stage GWA scenario employed in pQTL studies [4, 8, 9]. Namely, we contrast the results of LOCUS with those of GEMMA applied to the Ottawa and DiOGenes data, but this time considering all 4M SNPs available before LD pruning and employing a standard genome-wide Bonferroni correction of *α* = 0.05. We acknowledge that the 0.05-Bonferroni correction may be overly conservative and also discuss the results using a permissive 0.2-Bonferroni threshold (Material and methods).

The number of locus-protein hits identified and replicated with at least one of the two methods was 100, using Bonferroni correction of 0.05 for GEMMA (full list of SNP-protein hits in S5 Table and summarized by sentinel SNPs in S6 Table). Among these 100 hits, GEMMA missed 15 hits validated by LOCUS (9 with the permissive Bonferroni correction of 0.2). As many as 5 of these 15 hits were *trans* associations and 6/15 hits were not previously described in the literature using a *r*^2^ > 0.5-proxy search (Fig 2a and S5 Table); these two observations highlight again the ability of LOCUS to identify weak/*trans* effects that may go unnoticed by univariate analysis, in line with the above simulation studies. Moreover, all 4 hits whose protein was quantified by both proteomic technologies had a successful dual replication using the alternative technology.

Inspecting the estimates for the loci missed by GEMMA and validated by the LOCUS two-stage analyses indicates that some univariate signal is present but is too weak to be detected after multiplicity correction. In contrast, the multiplicity-adjusted LOCUS analyses could effectively single out and validate individual hits among the strongest GEMMA signals. The S3 Appendix provides regional association plots for six such examples: three loci associated with the CO7, ECM1 and ITIH3 MS levels, and three loci associated with the I17RB, PROC and TENA SomaLogic levels. Moreover, the superiority of LOCUS appears to be robust to the *p*-value threshold used for the univariate analysis: the replication rates and number of loci validated by GEMMA univariate design for a range of thresholds are inferior to those of LOCUS multivariate design (S3 Appendix).

Finally, LOCUS missed 6 locus-protein hits identified by GEMMA, but, importantly, these hits were not detectable by LOCUS. Indeed, all 115 SNPs selected by GEMMA had been removed by the *r*^2^ > 0.95 pruning employed to define the tag SNP panel used for the LOCUS study. To assess whether LOCUS would have identified these hits were their corresponding loci included in the primary analysis, we reran our method after adding to our original panel all 115 SNPs and their neighbours (1Mb span). LOCUS could successfully identify all 6 locus-protein hits with very high posterior probabilities of inclusion (PPI > 0.99). The sentinel SNPs selected by LOCUS are reported in the “LOCUS extended panel” column of S6 Table.

In summary, the improvement seen by simulation of the multivariate design over the classical univariate design is confirmed on the real data: LOCUS achieves higher replication rates, uncovers 15 pQTL loci missed by univariate screening (and does not miss any univariate signal when the corresponding SNPs are properly included in the tag-SNP panel). Among the hits missed by the univariate design but validated by LOCUS two-stage analysis, several are *trans* signals (proteins: INSR, PROC, SEM3A, TENA, XRCC6, i.e., 5/15), novel signals (CYTN, CYTT, FA12, FCN3, ITIH3, XRCC6, i.e., 6/15) and have successful cross-assay validation using the MS and SomaLogic measurements of their protein if available (CFAB, CO7, ECM1, XRCC6, i.e., 4/4).

### Two examples: a pleiotropic locus and a *trans*-acting locus

In this section we examine the advantages of our multivariate analysis for the detection of two loci, i.e., the notorious pleiotropic locus *ABO* and a novel locus on chromosome 11 *trans*-acting on the XRCC6 levels. These two examples are representative of the types of signals which LOCUS tends to better detect, namely weaker signals, possibly involving a shared control from a single locus. We also formulate some hypotheses on the involvement of these pQTLs in metabolic health, to be explored in dedicated studies.

The recovery of the *ABO* signals by LOCUS was facilitated by its tailored parametrization that directly models pleiotropy via *ω_s_*, as discussed in the simulation section and shown in Fig 3e. Fig 4a indicates that LOCUS piled up evidence on the two sentinels SNPs rs2519093 and rs8176741. Althougth the functional relevance of these two SNPs is by no means guaranteed and would need to be inspected in proper wet-lab or fine-mapping experiments, LOCUS could capitalize on the pleiotropic potential of these SNPs to effectively exploit the shared associations across the five proteins. Presumably because GEMMA does not leverage joint patterns across proteins, the univariate estimates failed to reach significance for the INSR protein, even using a loose Bonferroni threshold of α = 0.2.

**Figure 4:**
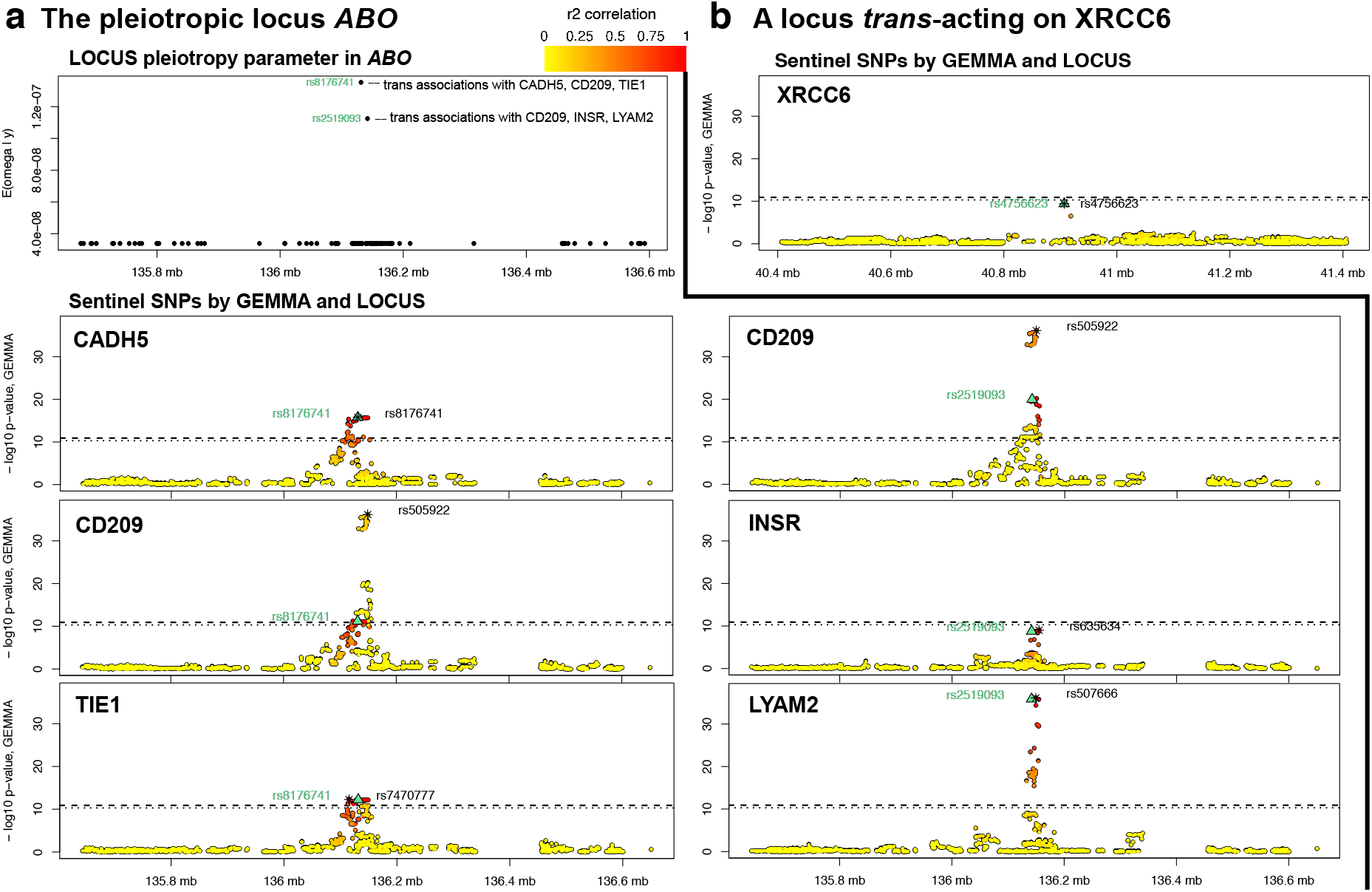
Two examples of pQTL signal estimation in the discovery set. (a) The pleiotropic locus *ABO*: The panel “LOCUS pleiotropy parameter in ABO” shows the posterior mean E (*ω_s_* | *y*) for a 1 Mb region around the gene ABO. This quantity attributes weight to two SNPs, rs8176741 and rs2519093, which LOCUS finds associated with the proteins CADH5, CD209, TIE1, resp., CD209, INSR, LYAM2. For each of these proteins, the colored panel displays the – log_10_ nominal *p*-values obtained when re-analyzing the Ottawa data with GEMMA [25, 26]; the dashed and dotted horizontal lines show the Bonferroni level of *α* = 0.05, resp. *α* = 0.2. The pleiotropic SNP identified by LOCUS is marked with a green triangle (rs8176741 left column, rs2519093 right column), and its correlation in *r*^2^ with the surrounding SNPs is indicated by the yellow to red colors. The top SNP found by GEMMA is shown by a black star; the univariate signal for INSR does not pass the *p*-value significance thresholds yet is detected by LOCUS multivariate analyses. (b) A locus *trans*-acting on XRCC6 (same labelling as in (a)).

*ABO* is a well-known pleiotropic locus associated with coronary artery diseases, type 2 diabetes, liver enzyme levels (alkaline phosphatase) and lipid levels [4, 5, 7]. Our analyses highlighted two independent sentinel SNPs in the *ABO* region: rs2519093 and rs8176741 (*r*^2^ = 0.03). The former SNP is *trans*-acting on E-selectin (protein LYAM2 encoded by SELE), the Insulin Receptor and the CD209 antigen. The latter SNP is *trans*-acting on the Tyrosine-protein kinase receptor (Tie-1), Cadherin-5 and CD209. Both SNPs were reported as cis-acting eQTL variants for *ABO, OBP2B* and *SURF1*, and further queries in public databases indicated that rs8176741 may affect the binding sites for three transcription factors (Myc, MYC-MAX and Arnt), suggesting a complex gene regulation circuitry.

Our analyses of the clinical parameters measured in the Ottawa and DiOGenes studies found significant associations of the controlled proteins with several clinical variables, such as triglycerides, visceral fat, fasting insulin, fasting glucose and HDL (Fig 5); this suggests that the *ABO* locus might have a role in metabolic health, although further research is needed to confirm this. In particular, the CD209 protein could be a secreted protein, released by M2 macrophages from adipose tissue, with a beneficial role in controlling lipid levels, thereby possibly protecting from developing dyslipidemia and related metabolic complications; see S4 Appendix for further discussion.

**Figure 5:**
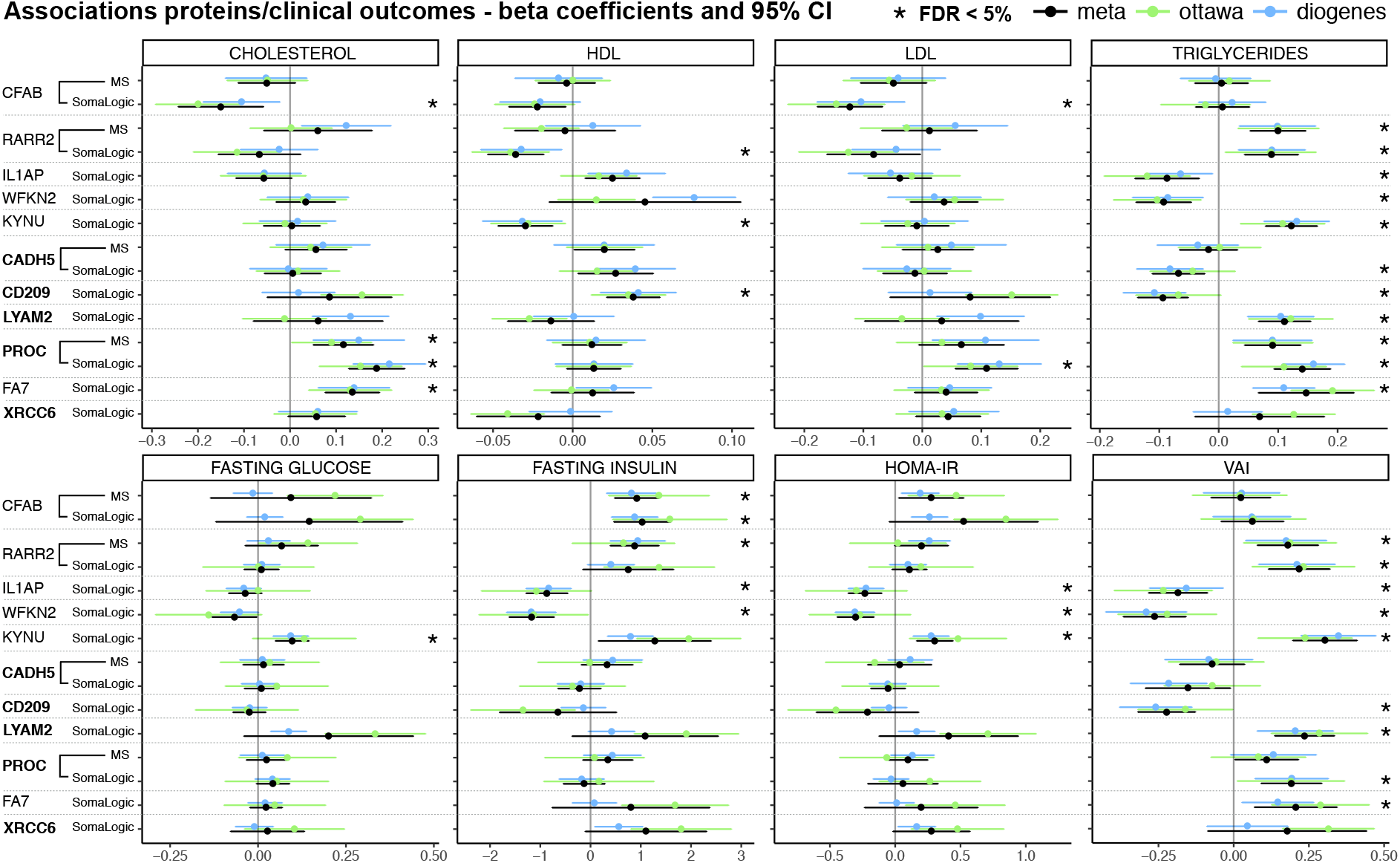
Forest plots for associations between proteins under genetic control and clinical parameters. Analyses were adjusted for age, gender and BMI (Material and methods) and the selection of proteins shown covers the *ABO* and XRCC6 pQTLs discussed in the main text as well as other examples discussed in S4 Appendix. All endpoints are measured in both the Ottawa a nd DiOGenes cohorts; they correspond to total lipid levels (first row: total cholesterol, HDL, LDL, triglycerides), glucose/insulin resistance (second row: fasting glucose, fasting insulin, HOMA-IR) and the visceral adiposity index (VAI). In each case, regression coefficients with 95% confidence intervals are shown for the Ottawa and DiOGenes analyses, and for the meta-analysis. The stars indicate associations with meta-analysis FDR < 5% (correction applied across all proteins under genetic control, not only those displayed). For proteins with measurements in the MS and SomaLogic platforms, association results are displayed for both; trans-regulated proteins are in bold.

Finally, some proteins had been reported as associated with the *ABO* locus [4] and yet did not pop up in our analyses (nor in the GEMMA analyses), namely, BCAM, LYAM3 (*SELP*), MET, NOTC1, OX2G (*CD200*), TIE2 (*TEK*), VGFR2 (*KDR)*, VGFR3 (*FLT4*), VWF. Multiple reasons could explain this lack of significance, including but not only: the use of samples from the obese population where these signals may be absent, a lack of statistical power and/or a poor quality of the SomaLogic measurements. For TIE2 and VGFR3, LOCUS reported very weak signals (PPI < 0.1) suggesting insufficient power. As for the other absent signals, the hypothesis of poorly quantified SomaLogic levels is not unlikely but difficult to assess since all seven proteins were measured only in the SomaLogic assay only and could therefore not be compared to corresponding MS measurements.

Our second example concerns the variant rs4756623 identified by LOCUS as a novel *trans* pQTL for the XRCC6 levels (X-Ray Repair Complementing Defective Repair In Chinese Hamster Cells; also known as Ku70). Proxy searches (down to *r*^2^ = 0.5 in European ancestry panels) did not reveal any tag SNP previously reported as a QTL (including e-, p-, or m-QTL). Again, the univariate screening results indicated some *trans* signal between rs4756623 and XRCC6, but the effect did not survive the multiplicity correction (Fig 4b).

The *XRCC6* gene activates DNA-dependent protein kinases (DNA-PK) to repair double-stranded DNA breaks by nonhomologous end joining. DNA-PKs have been linked to lipogenesis in response to feeding and insulin signaling [29]. DNA-PK inhibitors may reduce the risk of obesity and type 2 diabetes by activating multiple AMPK targets [30]. A recent review discussed the role of DNA-PK in energy metabolism, and in particular, the conversion of carbohydrates into fatty acids in the liver, in response to insulin [31]. It described increased DNA-PK activity with age, and links with mitochondrial loss in skeletal muscle and weight gain. Finally, *XRCC6* functions have been reported as associated with regulation of beta-cell proliferation, islet expansion, increased insulin levels and decreased glucose levels [30, 32].

We found that XRCC6 was significantly associated with decreased HDL, increased triglycerides, insulin levels and visceral adiposity in our studies, while rs4756623 is located within the *LRRC4C* gene a binding partner for Netrin G1, which has a known role in metabolism (S4 Appendix); the regulatory mechanisms between the rs4756623 locus and the XRCC6 levels should be clarified, and functional studies will be required to understand their physiological impact.

The next three sections provide a general characterization of the pQTLs identified by LOCUS in terms of their functional enrichment, overlapping patterns with known loci and clinical relevance of their associated proteins.

### Overlap with eQTLs and evidence for regulatory impact

We assessed the overlap of the 113 validated pQTLs with known eQTLs (S7 Table). Seventy-seven of the 104 sentinel SNPs involved in our pQTL associations had one or more eQTL associations in at least one tissue. These SNPs have been implicated in 83 eQTL associations, representing a significant enrichment (*p* < 2.2 × 10^−16^). Forty-nine of these 77 SNPs were eQTL variants for the gene coding for the protein with which they were associated in our datasets. Our pQTLs were also enriched in epigenome annotation marks (*p* = 9.20 × 10^−4^) and significantly closer to transcription start sites compared to randomly chosen SNP sets (*p* = 9.99 × 10^−6^). These observations suggest potential functional consequences for our pQTL hits.

### Overlap with GWAS risk loci

A total of 217 previously reported genome-wide associations overlapped our validated pQTL loci, corresponding to 139 unique traits mapping to 68 distinct regions (based on LD *r*^2^ > 0.8). Nineteen SNPs were directly involved in these associations (S8 Table) representing a significant enrichment (*p* < 2.2 × 10^−16^).

This suggests that studying the genetic associations with the proteome may serve as starting point to elucidate the molecular processes between the genotype and certain clinical endpoints whose genetic control has been confirmed, and acquire mechanistic insights on the pathways involved. Colocalization analyses are first step towards this end. The next section also paves the way for a discussion on the clinical relevance of our pQTL results by studying associations with metabolic parameters directly measured in the Ottawa and DiOGenes studies.

### Proteins as endophenotypes to study the genetic architecture of clinical traits

As briefly illustrated in the discussion of the XRCC6 *trans* effect and the *ABO* pleiotropy, the large panel of clinical variables pertaining to metabolic health measured in both the Ottawa and DiOGenes studies is a rich resource for evaluating the possible biomedical value of the validated pQTLs.

We performed a meta-analysis of the DiOGenes and Ottawa clinical and proteomic data, and found that 35 of the 88 proteins under genetic control had associations with dyslipidemia, insulin resistance or visceral fat-related measurements at FDR 5%. These associations should be attributable metabolic factors independently of overall adiposity, as we controlled for BMI as a potential confounder; they are listed and displayed as a network in S4 Appendix. We observed consistent directions of effects in the two cohorts (see examples of Fig 5, and S9 Table for full details). Based on these associations, S4 Appendix expands on the possible biomedical relevance of a selection of pQTLs in the context of metabolic complications.

Importantly, the set of 88 genetically-driven proteins was significantly more associated with the clinical variables compared to protein sets chosen randomly among all proteins quantified by our MS and SomaLogic panels (*p* = 0.014), with this background adjustment accounting for the bias of SomaLogic proteins towards specific inflammation and oncology-related pathways (S10 Table). This enrichment therefore supports that the primary pQTL analyses can help uncover potential proteomic biomarkers for the Metabolic Syndrome and related pathologies; the full pQTL results of S4 Table and meta-analysis clinical associations of S9 Table constitute a useful resource towards this end.

### Discussion

Despite important technological advances, large-scale pQTL studies remain infrequent, owing to their high costs [4–9]. To date, due to a lack of scalable multivariate methods, current QTL analyses rely on single-SNP/single-protein mapping, which inherently limits their findings to loci displaying strong effect sizes.

Here, we described the first integrative pQTL study that analyzes jointly all SNPs and proteomic levels from complementary MS and aptamer-based assays. Our Bayesian approach LOCUS confirmed 93 pQTLs (75 distinct proteins) highlighted in previous studies [4, 6–9], despite our sample sizes 2.5 to 18 times smaller, and revealed 20 novel pQTLs (18 distinct proteins, S4 Table) with sound evidence for functional relevance and possible implications in the development of the Metabolic Syndrome. Our two-stage analysis achieved very high replication rates (83%), and appreciable cross-technology validation for the pQTLs whose proteins had dual-assay measurements (78%). Importantly, 15% of the validated findings would have been missed by a standard univariate design.

This corroborates our extensive simulation studies, which demonstrated the improved QTL detection of LOCUS compared to univariate approaches on synthetic data mimicking the real data. In particular, these experiments indicated that the joint modelling of all outcomes leads to a significant increase of power in scenarios where a shared regulatory architecture governs groups of molecular levels. They also showed that accounting for local linkage disequilibrium using simulated annealing further enhances the detection of weak effects and drastically reduces the multiplicity burden. These advantages were exemplified for the pleiotropic locus ABO and a novel locus *trans*-acting on XRCC6, and can be particularly important on datasets with small sample sizes.

Finally, our analyses indicated that proteins under genetic control are enriched in associations with metabolic parameters, which supports a genetic basis of these parameters and emphasizes the advantages of pQTL studies for elucidating their underlying functional mechanisms. Evaluating the genetic control of plasma protein levels has the potential to reveal circulating biomarkers of direct interest for diagnostic applications but clearly, large pQTL studies in adipose tissues or liver may provide further, more specific insights on the role of the proteome in metabolic health. Although still at their early stages, pQTL studies are also intrinsically limited by the invasive nature of accessing tissues in humans. We hope that future progress in the availability of tissue-specific proteomic samples will reveal new mechanisms unavailable from plasma studies. As it is likely that sample sizes will remain limited, our well-powered joint approach will be a useful tool to this end. In the meantime, our complete plasma pQTL and clinical association results offer opportunities to study functional mechanisms and possible therapeutic options; they are accessible from the searchable online database https://locus-pqtl.epfl.ch/db.

Our work presents a direct illustration of a fully joint QTL analysis at scale and highlights concrete biological findings that take advantage of our tailored statistical approach. A central ambition was to showcase that LOCUS can bridge the gap between Bayesian multivariate inference and its practical use for analyzing current molecular QTL data. Indeed, the applicability of LOCUS goes beyond pQTL studies, as it is tailored to any genomic, proteomic, lipidomic, metabolic or methylation QTL analyses and can be used for genome-wide association with several clinical endpoints. The computational burden involved in the most Bayesian inference approaches has hampered the use of multivariate hierarchical modelling on large molecular QTL datasets thus far; our joint framework overcomes this with a scalable batch-wise variational algorithm and an effective C++/R implementation. It therefore offers a direct alternative to univariate screening approaches whose drawbacks for uncovering weak QTL effects are the object of a broad consensus [11–13]. Performance profiling for LOCUS demonstrated that it can analyze millions of SNPs and thousands of molecular levels (S2 Appendix). To our knowledge, no other fully multivariate method is applicable to large molecular QTL studies without drastic preliminary dimension reduction; our pilot study therefore opens new perspectives for uncovering weak and complex effects.

## Material and methods

### Ethics approval and consent to participate

The *Ottawa* and *DiOGenes* studies were approved by the local human research ethic committees. Participants provided informed written consent, and all procedures were conducted in accordance with the Declaration of Helsinki. Trial registration number: NCT00390637. Registered 20 October 2006.

### Study samples

The Ottawa study was a medically supervised program set up by the Weight Management Clinic of Ottawa [15]. Subjects under medication known to affect weight, glucose homeostasis or thyroid indices were excluded from all analyses.

The DiOGenes study was a multi-center pan-European program [16]. Eight partner states participated to the study: Bulgaria, the Czech Republic, Denmark, Germany, Greece, the Netherlands, Spain and the United Kingdom. Participants were overweight/obese (BMI between 27 and 45 kg/m^2^), non-diabetic and otherwise healthy.

For both studies, subjects who were not under fasting conditions at plasma sample collection were excluded from the proteomic analyses. The main clinical characteristics of the cohorts are given in S1 Table.

### Proteomic data

Plasma protein expression data were obtained using two types of technologies: mass-spectrometry (MS) and a multiplexed aptamer-based assay developed by SomaLogic [17]. Samples were randomized, ensuring that the plate numbers were not associated with age, gender, ethnicity, weight-related measures, glycemic indices, measures of chemical biochemistry, and, for the DiOGenes samples, collection centers.

The MS proteomic quantification used plasma samples spiked with protein standard lactoglobulin (LACB). Samples were immuno-depleted, reduced, digested, isobarically 6-plex labeled and purified. They were analyzed in duplicates on two separate but identical systems using linear ion trap with Orbitrap Elite analyzer and Ultimate 3000 RSLCnano System (Thermo Scientific). Protein identification was done with the UniProtKB/Swiss-Prot database [33], using Mascot 2.4.0 (Matrix Sciences) and Scaffold 4.2.1 (Proteome Software). Both peptide and protein false discovery rates (FDR) were set to 1%, with a criterion of two unique peptides. The relative quantitative protein values corresponded to the log_2_-transformation of the protein ratio fold changes with respect to their measurements in the biological plasma reference sample. The sample preparation and all other manipulations relative to the MS measurements are detailed further in previous work [34–36].

The SomaLogic protein measurements were characterized using the SOMAscan assay [17], which relies on fluorescent labelling of poly-nucleotide aptamers targeting specific protein epitopes. Protein measurements were obtained in relative fluorescence unit and were then log_2_-transformed. The SomaLogic panel targetted 1, 129 proteins; the KEGG and Reactome pathways covered by this panel are listed in S10 Table (enrichment using a 5% FDR threshold).

We discarded MS-based proteins if their measurements were missing for more than 5% of the samples, leaving 210 proteins in the Ottawa cohort and 136 in the DiOGenes cohort; we restricted all downstream analyses to the 133 proteins available for both cohorts. The SomaLogic measurements had no missing values. Totals of 1, 100 and 1, 129 proteins were assayed in the Ottawa and DiOGenes cohorts. All our analyses focused on the 1, 096 proteins quantified for both cohorts. The overlap between the MS and SomaLogic panels was of 72 proteins for 397 and 389 subjects in Ottawa and DiOGenes respectively.

We excluded samples with extreme expression values in more than 5% of the proteins, i.e., values beyond the outer fences of the empirical distribution (*q*_1_ – 3 × IQR, *q_3_* + 3 × IQR, where *q*_1_, *q*_3_ are the lower and upper quartiles, and IQR is the interquartile range). After this quality control procedure, 577 and 428 Ottawa samples remained in the MS and SomaLogic datasets, respectively, and 481 and 563 DiOGenes samples remained in the MS and SomaLogic datasets, respectively.

### Genotyping

Genotypes were generated using HumanCoreExome-12 v1.1 Illumina SNP arrays (Illumina, Inc., San Diego, CA), according to their manufacturer’s instructions and were called with the GenomeStudio Software provided by Illumina. Preprocessing steps, including imputation and quality control, have been previously documented [37]. We discarded SNPs with call rate < 95%, violating Hardy–Weinberg equilibrium (FDR < 20%), and we discarded subjects with low call rate (< 95%), abnormally high autosomal heterozygosity (FDR < 1%), an XXY karyotype, or gender inconsistencies between genotype data and clinical records. For subjects with identity-by-state IBS> 95%, we kept only the one with the highest call rate. A table detailing these quality control filters for the Ottawa and DiOGenes studies is given in S5 Appendix. The subjects from both cohorts were of European ancestry and the two cohorts had similar genetic structure. We used principal component analyses separately on each cohort to exclude subjects that were extremely heterogeneous genetically. We performed genotype imputation using SHAPEIT [38] and IMPUTE2 [39], based on the European reference panel from the 1, 000 Genome project (March 2012 release, phase 1 version 3). We then discarded SNPs with INFO score < 0.8, which left 4.9M imputed SNPs in both datasets. We applied a light LD pruning with PLINK [40] using pairwise *r*^2^ threshold 0.95 and used a minor allele frequency threshold of 5% after having restricted the genotype data to the subjects with available proteomic data.

The above steps were performed separately for the Ottawa and the DiOGenes cohorts, so in order to define a common set of SNPs for discovery and replication, we restricted each dataset to the SNPs available for both cohorts. After all genetic quality controls, and in both cohorts, *p* = 275,485 tag SNPs remained for the SomaLogic analysis and *p* = 275, 297 tag SNPs remained for the MS analysis. As SNPs were imputed, the 0.95-*r*^2^ pruning led to a drastic cut of “redundant” markers: without this pruning step, the number of SNPs was ≈ 4M. Such a reduction is not surprising considering the nature of the underlying SNP arrays (essentially based on tag SNPs) and indicates that little information was discarded. In the Ottawa cohort *n* = 376 subjects had both genotype and MS proteomic data, and *n* = 394 subjects had both genotype and SomaLogic proteomic data. In the DiOGenes cohort, these numbers were *n* = 400 and 548.

### Clinical data

Both cohorts had records on age, gender, anthropometric traits (weight and BMI), glycemic variables (fasting glucose, fasting insulin, HOMA-IR), and total lipid levels obtained from blood biochemistry (total cholesterol, triglycerides, HDL). We derived LDL values using the Friedewald formula [41], and obtained gender-specific *visceral adiposity index* (VAI) values using the formula of Amato et al. [42]. In each cohort and for each clinical variable, we removed a few samples with extreme measurements, similarly as for the proteomic data quality control.

### LOCUS: fast Bayesian inference for joint QTL analysis

LOCUS is a variational inference approach for joint mapping analysis at the scale required by current molecular QTL studies (Fig 1a) [14]. It implements a hierarchical model that involves a collection of sparse regressions,

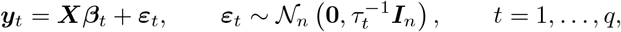

where ***y*** = (***y***_1_,…, ***y***_q_) is an *n × q* matrix of *q* centered outcomes (e.g., genomic, proteomic or metabolomic levels), and ***X*** is an *n × p* matrix of *p* centered candidate predictor SNPs, for each of *n* samples. Each outcome, ***y**_t_*, is related linearly to all *p* candidate SNPs, and has a specific residual precision, *τ_t_*, to which we assign a Gamma prior, *τ_t_* ~ Gamma(*η_t_, κ_t_*). The association between each pair of SNP and outcome is modelled using a spike-and-slab prior, namely, for *s* = 1,…, *p* and *t* = 1,…, *q*,

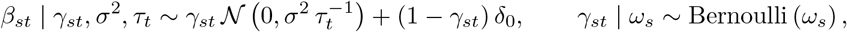

where δ_0_ is the Dirac distribution. Hence, to each regression parameter *β_st_* corresponds a binary latent parameter γ_*st*_, which acts as a “predictor-outcome association indicator”: the predictor **X**_s_ is associated with the outcome ***y**_t_* if and only if γ_*st*_ = 1. The parameter *σ* represents the typical size of nonzero effects and is modulated by the residual scale, 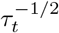, of the outcome concerned by the effect; it is inferred from the data using a Gamma prior specification, *σ*^−2^ ~ Gamma (λ,*ν*). Finally, we let the probability parameter *ω_s_* have a Beta distribution,

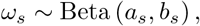

where *a_s_* and *b_s_* are set so as to enforce sparsity as described in Ruffieux et al. [14]. Since it is involved in the Bernoulli prior specification of all γ_*s*1_,…,γ_*sq*_, the parameter *ω_s_* controls the proportion of outcomes associated with the predictor ***X**_s_*, and hence directly represents the propensity of predictors to be pleiotropic “hotspots”. Both *ω_s_* and *σ*^2^ allow the leveraging of shared association patterns across all molecular variables, which enhances the estimation of QTL effects; see the graphical representation of the model (Fig 1b). Thanks to this joint tailored modelling, the inclusion of *cis* regions does not mask the weaker *trans* effects, but rather modelling altogether *cis* and *trans* effects, possibly governed by shared mechanisms, contributes to boost the detection of *trans* signals. LOCUS estimates interpretable posterior probabilities of association for all SNP-outcome pairs (Fig 1c), from which Bayesian false discovery rates are easily calculated.

The Bayesian framework of LOCUS can be flexibly extended to account for population structure, using a dedicated random effect parameter. Alternatively, adding genotypic principal components as fixed covariates in our model may suffice in some cases.

Inference on high-dimensional Bayesian models is both computationally and statistically difficult. Our previous joint QTL approaches [20, 43] are based on sampling procedures, such as MCMC algorithms, and require prohibitive computational times on data with more than few hundreds of SNPs or outcomes. LOCUS uses a fast deterministic variational inference algorithm, which scales to the typical sizes of QTL problems. Previous work [14] compared LOCUS with existing QTL methods, whether stochastic or deterministic, univariate or multivariate. Here, we augmented our algorithm with a simulated annealing procedure [44] to enhance exploration of multimodal parameter spaces, as induced by strong linkage-disequilibrium (LD) structures. The pQTL study of two obesity cohorts described in the present paper illustrates the strengths of our statistical framework in a concrete and thorough manner and directly exploits these strengths for uncovering new signals of biological and clinical relevance.

The applicability of a fully multivariate method to large molecular QTL data also hinges on the effective computational implementation of its algorithmic procedure. The annealed variational updates of LOCUS are analytical and performed by batches of variables. The software is written in R with C++ subroutines and is publicly available [45]. Our pQTL analyses completed in a few hours for 275K tag SNPs representing information from about 4M common markers, yet larger SNP panels can be considered as our method scales linearly in terms of memory and CPU usage. For instance, analyses of 2M SNPs and 1, 000 proteins run in less than 40 hours (see profiling, S2 Appendix).

### Simulation study design

We evaluated the performance of LOCUS expected on our data by conducting two simulation studies. We compared its statistical power to detect pQTL associations with that of the linear mixed model approach GEMMA [25, 26], which estimates the associations between each SNP and each outcome in a univariate fashion. We used the R package `echoseq` [46] to generate synthetic data that emulate real data; see S2 Appendix.

For the first simulation, we ran LOCUS and GEMMA on the SNPs of all *n* = 376 Ottawa subjects, and on simulated expression outcomes with residual dependence replicating that of the q = 133 MS proteomic levels. We used the SNPs from chromosome one (*p* = 20, 900), and generated associations between 20 SNPs and 25 proteins chosen randomly, leaving the remaining variables unassociated. Some proteins were under pleiotropic control; we drew the degree of pleiotropy of the 20 SNPs from a positively-skewed Beta distribution, so only a few SNPs were hotspots, i.e., were associated with many proteins. We generated associations under an additive dose-effect scheme and drew the proportions of outcome variance explained by a given SNP from a Beta(2, 5) distribution to give more weight to smaller effect sizes. We then rescaled these proportions so that the variance of each protein attributable to genetic variation was below 35%. These choices led to an inverse relationship between minor allele frequencies and effect sizes, which is to be expected under natural selection. We generated 50 replicates, re-drawing the protein expression levels and effect sizes for each. On top of comparing the relative performances of LOCUS and GEMMA on these data, we also illustrated the benefits of modelling all the proteomic outcomes jointly by comparing the performance of LOCUS when applied to increasingly larger batches of proteins. Namely, we split randomly the proteins into batches of given sizes – ranging from 1, for a univariate modelling of the proteins, to all 133 MS proteins, for a fully multivariate treatment corresponding to a classical application of LOCUS – and ran LOCUS separately on each batch and all SNPs from chromosome one. We then aggregated posterior probabilities of inclusion to assess selection performance. This procedure was repeated for each of the 50 synthetic data replicates.

For the second simulation, we re-assessed the performance of LOCUS for a grid of data generation scenarios. We considered a wide range of sparsity levels (numbers of proteins under genetic control) and effect sizes (proportions of outcome variance explained by the genetic variants). Given the large number of configurations (130), and in order to limit the computational burden, we used the first *p* = 2, 000 SNPs, and ran LOCUS and GEMMA on 20 replicates for each configuration.

### pQTL analyses

We performed pQTL analyses separately for each platform, i.e., one analysis for the MS proteomic dataset, and another for the SomaLogic proteomic dataset. Each analysis comprised two stages: a discovery stage using the Ottawa cohort and a replication stage based on the DiOGenes cohort.

For discovery, we used LOCUS on both the MS and the SomaLogic datasets, with an annealing schedule of 50 geometrically-spaced temperatures and initial temperature of 20; pilot experiments indicated that estimation was not sensitive to these choices. We used a convergence tolerance of 10^−3^ on the absolute changes in the objective function as the stopping criterion. The algorithm can handle missing data in the outcome matrix, so no imputation was necessary for the MS proteomic data.

We adjusted all analyses for age, gender, and BMI. No important stratification was observed in the genotype data; the first ten principal components together explained little of the total variance (< 4%), so we did not include them as covariates. We derived FDR values from the posterior probabilities of association obtained between each SNP and each protein, and reported pQTL associations using an FDR threshold of 5%.

We performed a validation study of the pQTLs discovered using the DiOGenes cohort with GEMMA [26, 47], with centered relatedness matrix (default) and *p*-values from (two-sided) Wald tests. We then obtained adjusted *p*-values using Benjamini–Hochberg FDR, and validated our hits using again an FDR threshold of 5%.

### Comparison with a standard two-stage univariate design

To assess the extent to which LOCUS two-stage pQTL analysis discovers more hits than the univariate procedures routinely applied for e- or pQTL analyses, we re-performed the entire study using GEMMA [25, 26]. We followed standard practices and ran the method separately for the MS and SomaLogic analyses on the SNPs without LD pruning, i.e., on roughly 4 million SNPs. We then corrected for multiple testing using a standard Bonferroni threshold of 0.05 (based on the numbers of tested SNPs and proteins) and also discussed the results obtained with a more permissive Bonferroni threshold of 0.2.

To account for proxy hits arising from the SNP LD structure and provide grounds for comparison between GEMMA and LOCUS, we defined hits at the level of loci as follows: the hits identified by GEMMA and/or LOCUS as associated with a same protein (quantified by the same proteomic technology) were considered to be in a same locus if there was no more than 1Mb between two consecutive hits. The additional hits found by GEMMA at Bonferroni level 0.2 were assigned to the closest existing loci (mapping to the same protein), provided that the distance was less than 1Mb; new loci were defined for the remaining hits.

### pQTL annotation

We used the Ensembl database [48] (GRCh37, release 94) to retrieve the list of genes within 2Mb of each *sentinel* SNP (i.e., involved in the pQTL associations identified by LOCUS), and also listed the SNPs in LD (*r*^2^ > 0.8), limiting the search to 500Kb upstream and downstream of the sentinel SNP position. We called *cis pQTLs*, all sentinel SNPs located within ±1Mb of the gene encoding for the controlled protein, and *trans pQTLs*, all other pQTLs.

We evaluated the overlap between our pQTL associations and previously reported pQTL signals with the PhenoScanner database [49, 50], using the default *p*-value threshold *p* < 10^−5^ and an LD proxy search (*r*^2^ > 0.8, in populations with European ancestry). As queries using the R-package `phenoscanner` are limited to 500 returned tuples, we downloaded a local copy of the database (retrieved on 26/03/2019). Moreover, since protein names in the database do not follow the official UniProt protein names, we retrieved the annotation files from all individual studies and remapped the protein names onto the official UniProt identifier, thereby enabling the comparison with our pQTL hits using dbSNP rsIDs and UniProt IDs.

### Epigenomic annotation

We retrieved epigenomic annotations of 1,000 Genomes Project (release 20110521) from Pickrell [51]. The data covered 450 annotation features, each binary-coded according to the presence or absence of overlap with the SNPs. The features corresponded to DNase-I hypersensitivity, chromatin state, SNP consequences (coding, non-coding, 5’UTR, 3’UTR, etc), synonymous and nonsynonymous status and histone modification marks. We obtained distances to the closest transcription start site from the UCSC genome browser [52]. Ninety-seven of our 104 validated sentinel SNPs had annotation data; to evaluate their functional enrichment, we resampled SNP sets of size 97 from our initial SNP panel, and, for each set, we computed the cumulated number of annotations. We did the same for the distances to transcription start sites. We repeated this 10^5^ times to derive empirical *p*-values.

### Overlap with known eQTLs and with GWAS risk loci

We evaluated the overlap of our pQTLs with the eQTL variants reported by the GTEx Consortium [53, 54] (release 7) at q-value < 0.05. We considered all 49 tissues listed by GTEx but eQTL SNPs for several tissues were counted only once. We made both general queries and queries asking whether a pQTL uncovered by LOCUS was an eQTL for the gene coding for the controlled protein.

We retrieved known associations between the validated sentinel pQTLs and diseases or clinical traits, based on the GWAS catalog [55, 56] (v1.0 release e92), and also using an LD proxy search (*r*^2^ > 0.8).

We evaluated enrichment for eQTL and risk loci using one-sided Fisher exact tests based on the 104 validated sentinel pQTLs.

### Associations with clinical variables

We tested associations between the proteins under genetic control and clinical parameters separately in each cohort. For the DiOGenes data, we used linear mixed-effect models, adjusting for age, gender as fixed effects, and center as a random effect. For the Ottawa data, we used linear models, adjusting for age and gender. Except when testing associations with anthropomorphic traits, all analyses were also adjusted for BMI. For the clinical variables available in the two cohorts (total cholesterol, HDL, LDL, fasting glucose, fasting insulin, HOMA-IR, triglycerides and VAI), we performed meta-analyses using the R package `metafor` [57]. We used random-effects models to account for inter-study variability, which may in part result from geographical differences, and employed two-sided Wald tests for fixed effects, and Cochran Q-tests for measuring residual heterogeneity; we did not interpret the results if between-study heterogeneity estimates were high (*I*^2^ > 80%), and evaluated the directional consistency of the effects between Ottawa and DiOGenes. We adjusted for multiplicity using Benjamini–Hochberg correction across all tests, i.e., involving the 88 tested proteins and the two proteomic technologies, and reported associations using a 5% FDR threshold.

We assessed whether the proteins under genetic control were enriched in associations with the clinical variables. We randomly selected 10^5^ sets of 88 proteins from the panel used for the pQTL analyses and derived an empirical *p*-value by counting, for each set, the number of proteins with at least one clinical association at FDR 5%.

## Supporting information

**S1 Appendix. Comparison of MS and SomaLogic measurements.** Scatterplots for the 72 protein levels with dual-technology quantification for both the Ottawa and the DiOGenes cohorts.

**S2 Appendix. Addendum to the simulation studies and computational performance of LOCUS.** Emulation of the MS QTL data; Runtime profiling for different problem sizes.

**S3 Appendix. Addendum to the comparison of LOCUS with the univariate analysis GEMMA.**

Six examples of pQTL loci found by LOCUS and missed by the univariate analysis; Sensitivity of the univariate analysis to the *p*-value threshold.

**S4 Appendix. Further examples of pQTL loci with possible implications in metabolic disorders.** Clinical associations with proteins controlled by the pleiotropic locus *ABO*. Clinical associations with the XRCC6 protein levels. CFAB and RARR2, mediators of adipogenesis are under genetic control; The importance of IL1AP for Metabolic Syndrome; WFKN2, a TGF*β*-activity protein with protective effect against metabolic disorders; Inflammation mediated proteins and their role in insulin resistance; Complement/coagulation: a *trans*-acting insertion linking PROC and its receptor.

**S5 Appendix. Addendum to the genotyping quality control.** Detailed description of the quality control performed on the genotyping data for the Ottawa and DiOGenes studies.

**S1 Table. Ottawa & DiOGenes cohorts.** Average anthropometric, glycemic and total lipid characteristics of the Ottawa and DiOGenes cohorts.

**S2 Table. Discovery pQTLs.** pQTL associations discovered by LOCUS in the Ottawa cohort, using the mass-spectrometry and SomaLogic datasets.

**S3 Table. Validation pQTLs.** pQTL associations discovered by LOCUS in the Ottawa cohort and validated in the DiOGenes cohort.

**S4 Table. Overlap with public pQTLs.** Overlap of the validated pQTL associations with public pQTL associations (PhenoScanner).

**S5 Table. Comparison with univariate analysis, individual SNP-protein pairs.** Comparison with the univariate two-stage analysis using GEMMA, individual hits.

**S6 Table. Comparison with univariate analysis, sentinel SNP-protein pairs.** Comparison with the univariate two-stage analysis using GEMMA, hits summarized by loci.

**S7 Table. Overlap with public eQTLs.** Overlap of the validated pQTL associations with public eQTL associations (GTEx Consortium).

**S8 Table. Overlap with risk loci.** Overlap of the validated pQTL associations with disease risk loci (GWAS Catalog).

**S9 Table. pQTLs vs clinical.** Associations between proteins under genetic control and clinical parameters measuring dyslipidemia, insulin resistance and visceral fat, in the Ottawa and DiOGenes cohorts.

**S10 Table. Pathways covered by the SomaLogic panel.** Enrichment analysis using KEGG (2019) and Reactome (2016).

## Availability of data and materials

The MS proteomic data have been deposited on the ProteomeXchange Consortium [58] via the PRIDE partner repository, with the dataset identifiers PXD005216 for DiOGenes and PXD009350 for Ottawa. The SomaLogic proteomic data are available from the Open Science Framework, at https://osf.io/v8mes/?view_only=13e4ccd127024ee7b4c819385325925c and https://osf.io/s4v8t/?view_only=90637f2941e14ec986e5888491fbdbbb, respectively for Ottawa and DiOGenes. All pQTL and clinical association results are available as supplemental tables and can be browsed from our online database https://locus-pqtl.epfl.ch/db. Other data that support the findings of this study are available from the corresponding authors upon request. All statistical analyses were performed using the R environment [59] (version 3.3.2). LOCUS and ECHOSEQ are freely available from GitHub [45, 46].

## Web ressources

ECHOSEQ: https://github.com/hruffieux/echoseq

Ensembl: http://grch37.ensembl.org/index.html

GEMMA: http://www.xzlab.org/software.html

GTEx: https://gtexportal.org/home

GWAS Catalog: https://www.ebi.ac.uk/gwas

IMPUTE2: http://mathgen.stats.ox.ac.uk/impute/impute_v2.html

JASPAR: http://jaspar.genereg.net

LOCUS: https://github.com/hruffieux/locus

Metafor: https://cran.r-project.org/web/packages/metafor/index.html

PhenoScanner: http://www.phenoscanner.medschl.cam.ac.uk

PLINK: http://zzz.bwh.harvard.edu/plink

ProteomeXchange: http://www.proteomexchange.org

R: https://www.r-project.org

SHAPEIT: https://mathgen.stats.ox.ac.uk/genetics_software/shapeit/shapeit.html

SNP2TFBS: https://ccg.vital-it.ch/cgi-bin/snp2tfbs/snpviewer_form_parser.cgi

UCSC: https://genome.ucsc.edu

UniProt: https://www.uniprot.org

## Acknowledgments

We thank Antonio Núñez Galindo, John Corthésy, Martin Kussmann, Ornella Cominetti and Loïc Dayon for designing and performing proteomic experiments, and the generation and cleaning of the MS-based proteomic datasets. We thank Zoltán Kutalik, Nele Gheldof and Ruth McPherson for their constructive comments, and thank Leonardo Bottolo and Sylvia Richardson for their contributions on the method’s simulated annealing procedure.

