## Supplementary material for "A fully joint Bayesian quantitative trait locus mapping of human protein abundance in plasma": S1 Appendix

### Proteins measured by the MS and SomaLogic technologies

Scatterplots for the 72 proteins having both MS and SomaLogic measurements, with loess fit. Blue plot titles indicate the proteins involved in validated pQTL hits.

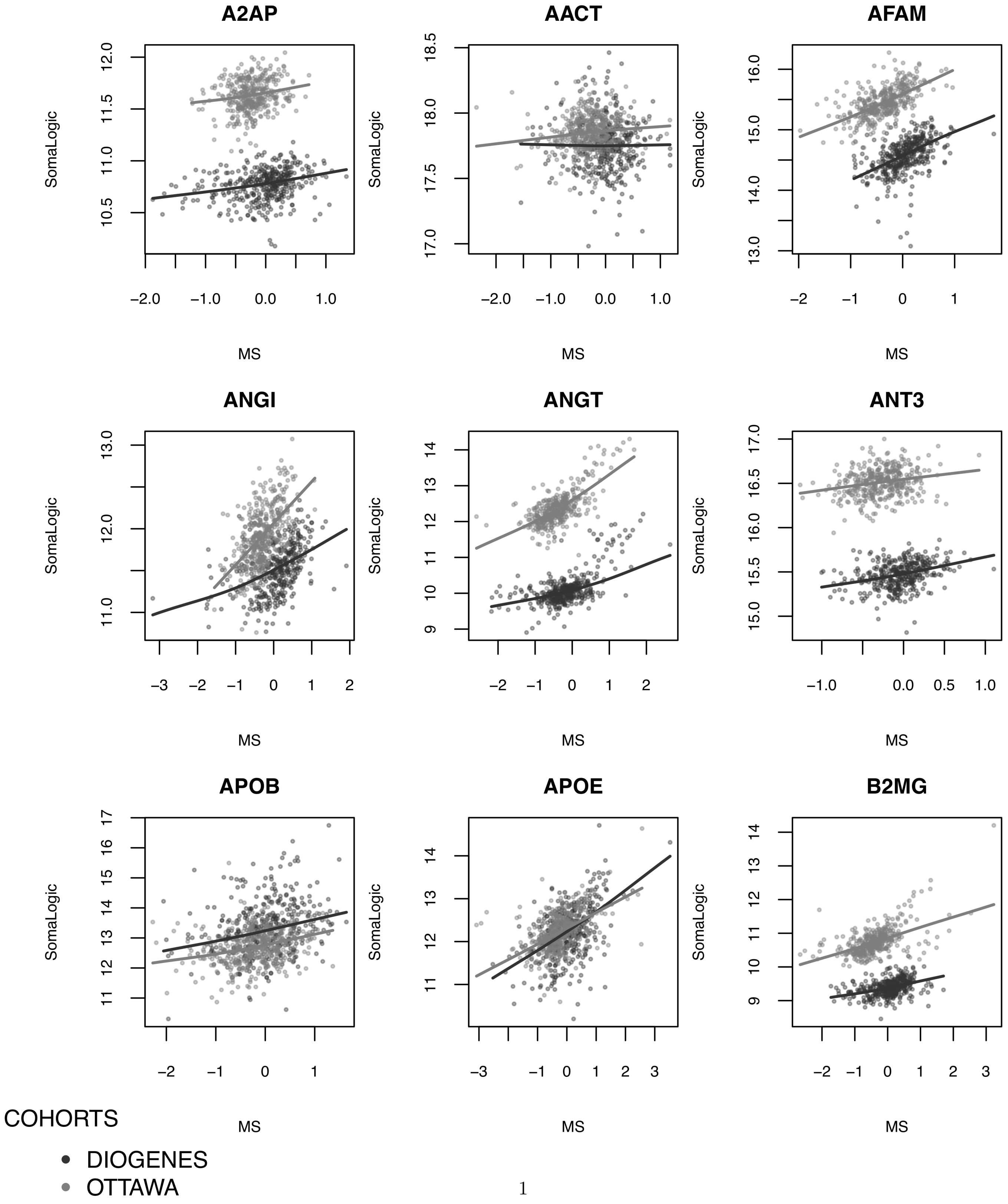

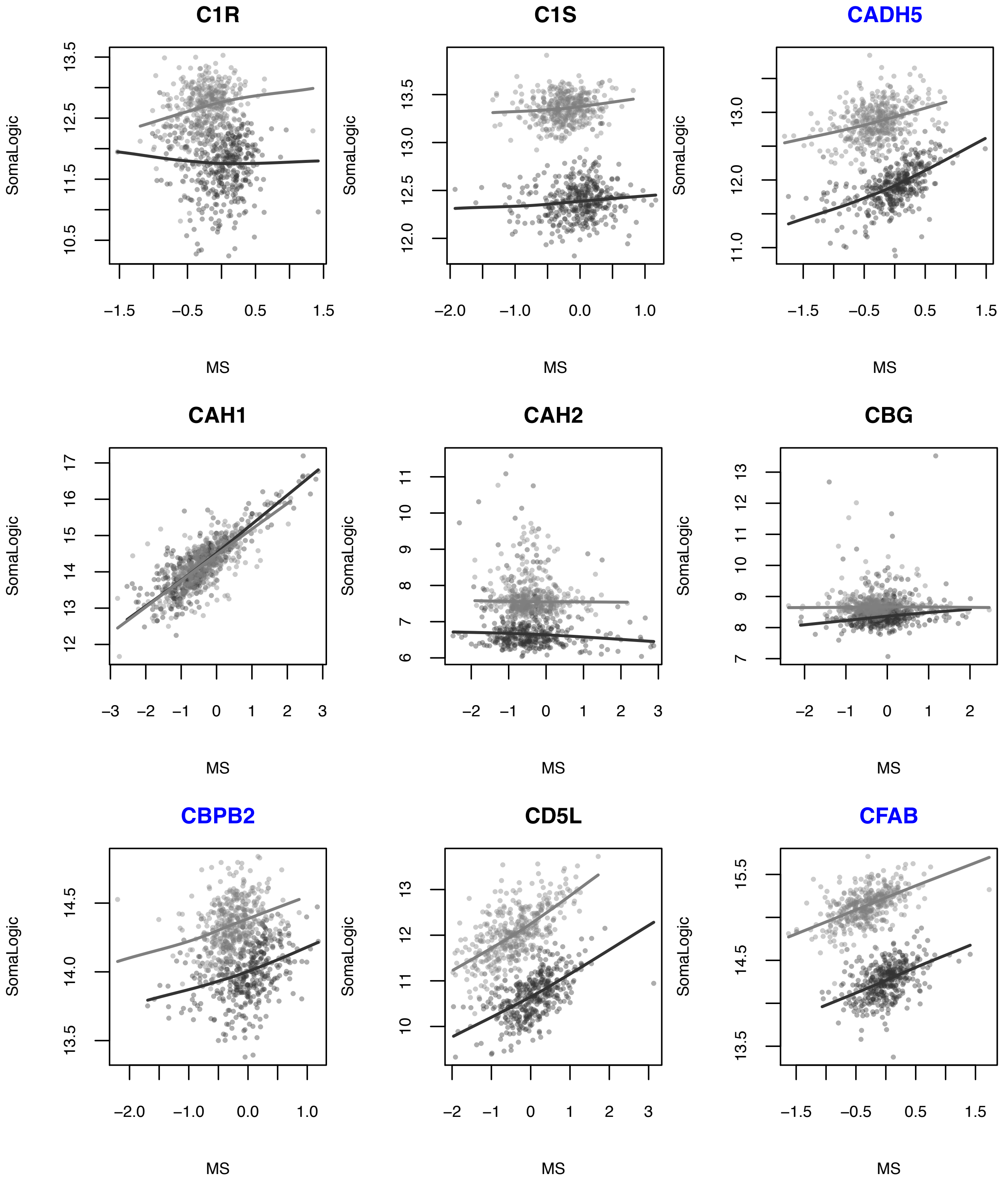
C1R
SomaLogic
MS
13.5
12.5
11.5
10.5
−1.5
−0.5
0.5
1.5
C1S
SomaLogic
MS
13.5
13.0
12.5
12.0
−2.0
−1.0
0.0
1.0
CADH5
SomaLogic
MS
13.0
12.0
11.0
−1.5
−0.5
0.5
1.5
CAH1
SomaLogic
MS
17
16
15
14
13
12
−3
−2
−1
0
1
2
3
CAH2
SomaLogic
MS
11
10
9
8
7
6
−2
−1
0
1
2
3
CBG
SomaLogic
MS
13
12
11
10
9
8
7
−2
−1
0
1
2
CBPB2
SomaLogic
MS
14.5
14.0
13.5
−2.0
−1.0
0.0
1.0
CD5L
SomaLogic
MS
13
12
11
10
−2
−1
0
1
2
3
CFAB
SomaLogic
MS
15.5
14.5
13.5
−1.5
−0.5
0.5
1.5

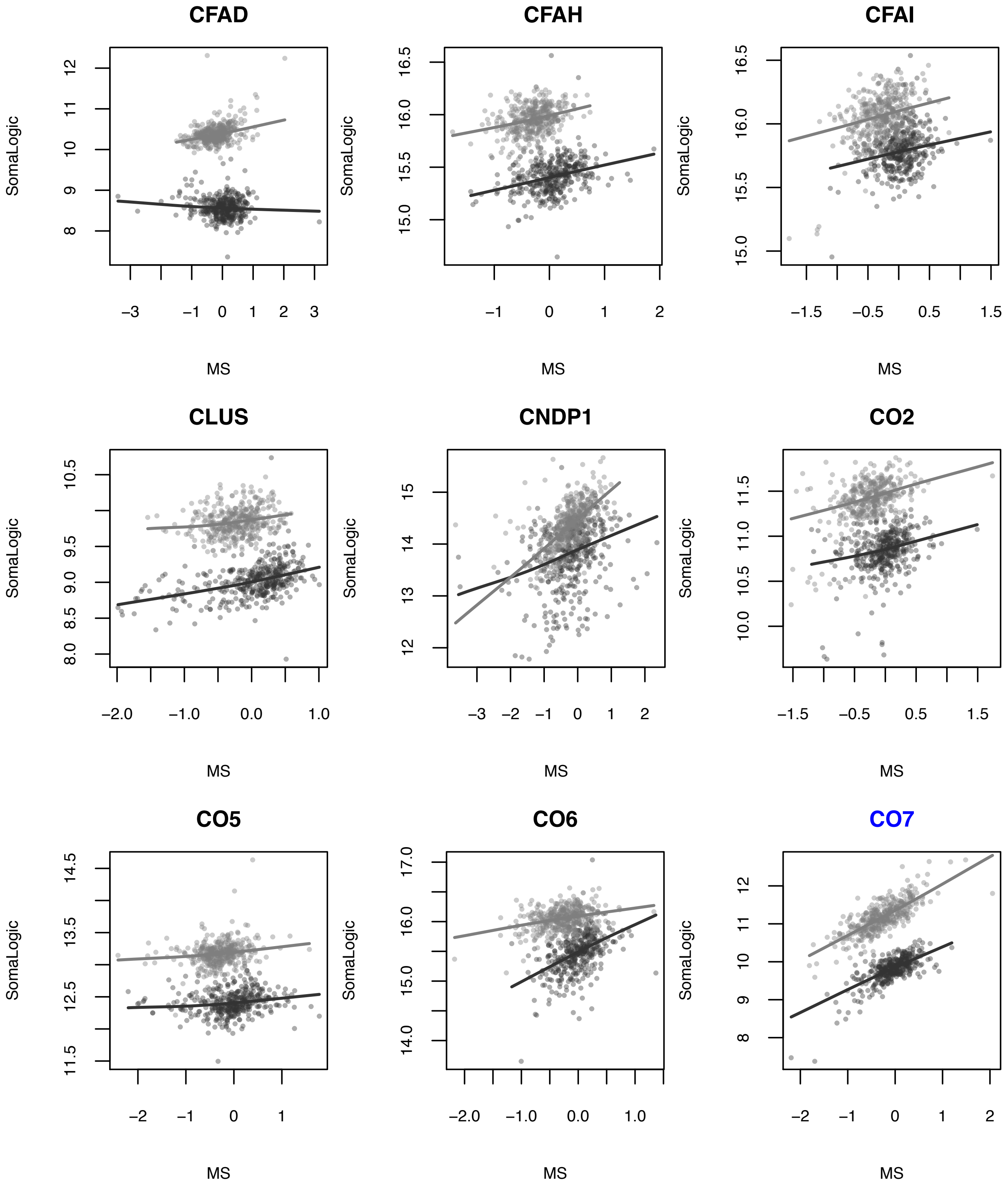
CFAD
CFAH
CFAI
CLUS
CNDP1
CO2
CO5
CO6
CO7
SomaLogic
MS

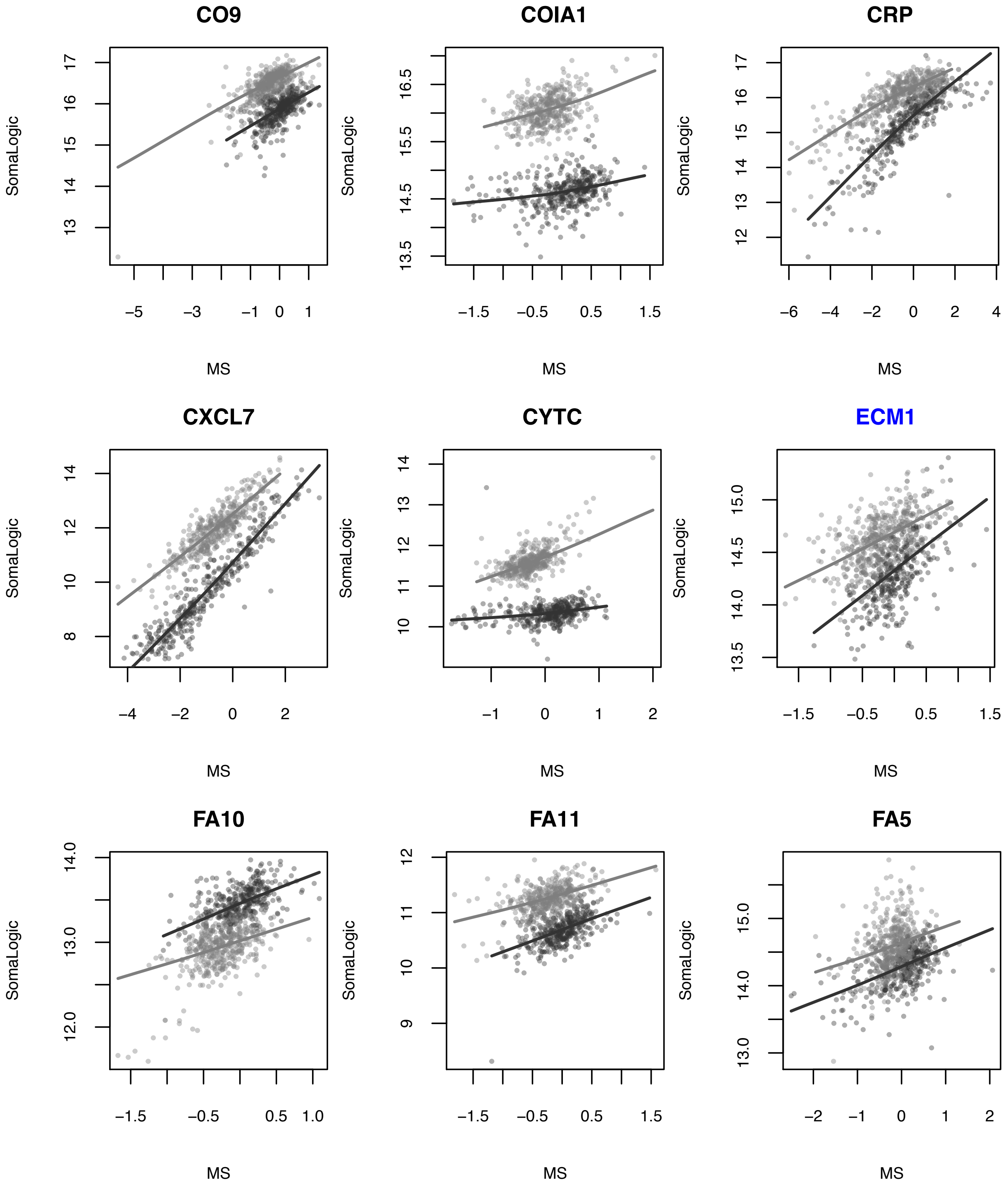
CO9
SomaLogic
MS
13
14
15
16
17
−5
−3
−1
0
1
COIA1
SomaLogic
MS
13.5
14.5
15.5
16.5
−1.5
−0.5
0.5
1.5
CRP
SomaLogic
MS
12
13
14
15
16
17
−6
−4
−2
0
2
4
CXCL7
SomaLogic
MS
8
10
12
14
−4
−2
0
2
CYTC
SomaLogic
MS
10
11
12
13
14
−1
0
1
2
ECM1
SomaLogic
MS
13.5
14.0
14.5
15.0
−1.5
−0.5
0.5
1.5
FA10
SomaLogic
MS
12.0
13.0
14.0
−1.5
−0.5
0.5
1.0
FA11
SomaLogic
MS
9
10
11
12
−1.5
−0.5
0.5
1.5
FA5
SomaLogic
MS
13.0
14.0
15.0
−2
−1
0
1
2

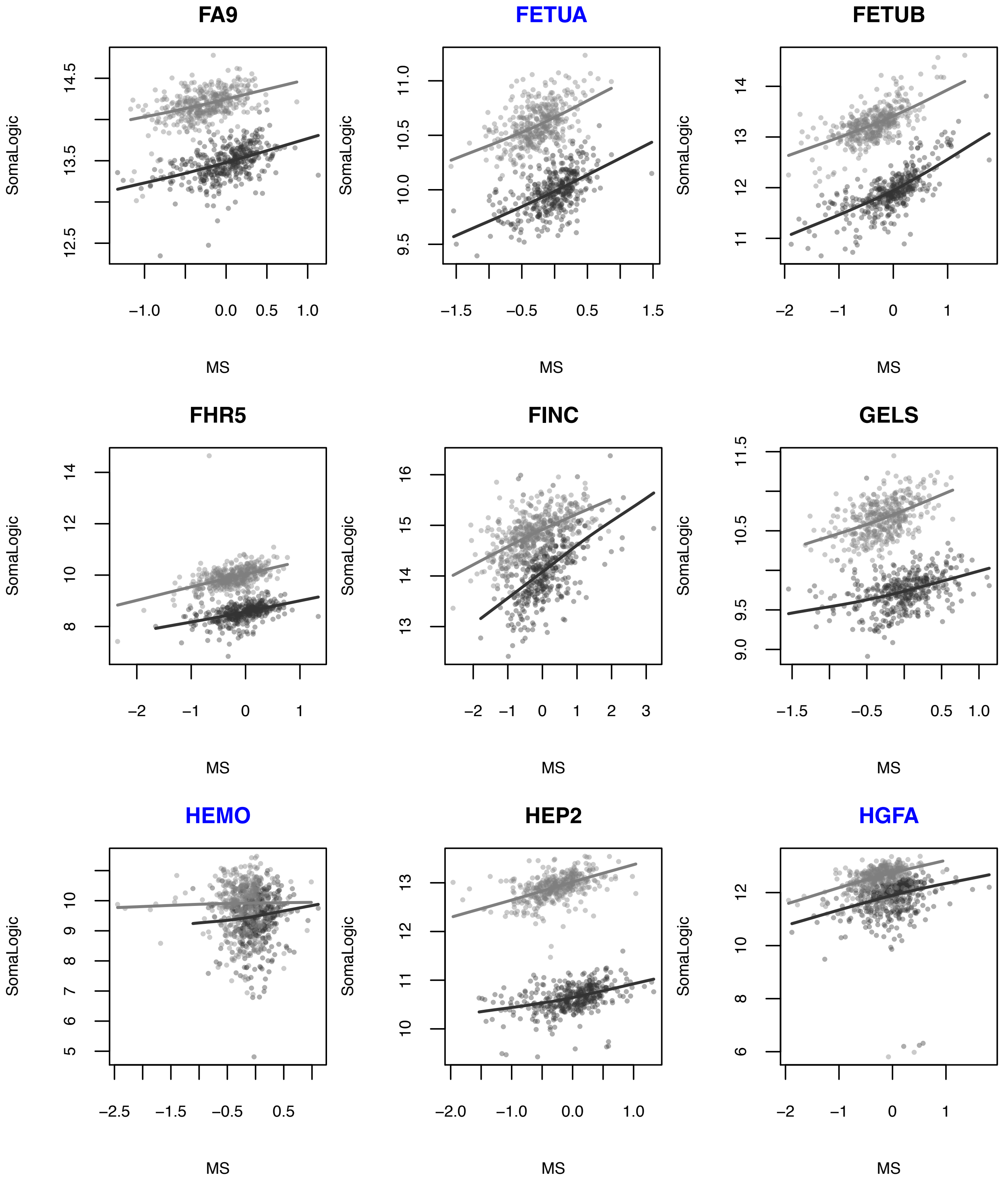
FA9
FETUA
FETUB
FHR5
FINC
GELS
HEMO
HEP2
HGFA
SomaLogic
MS

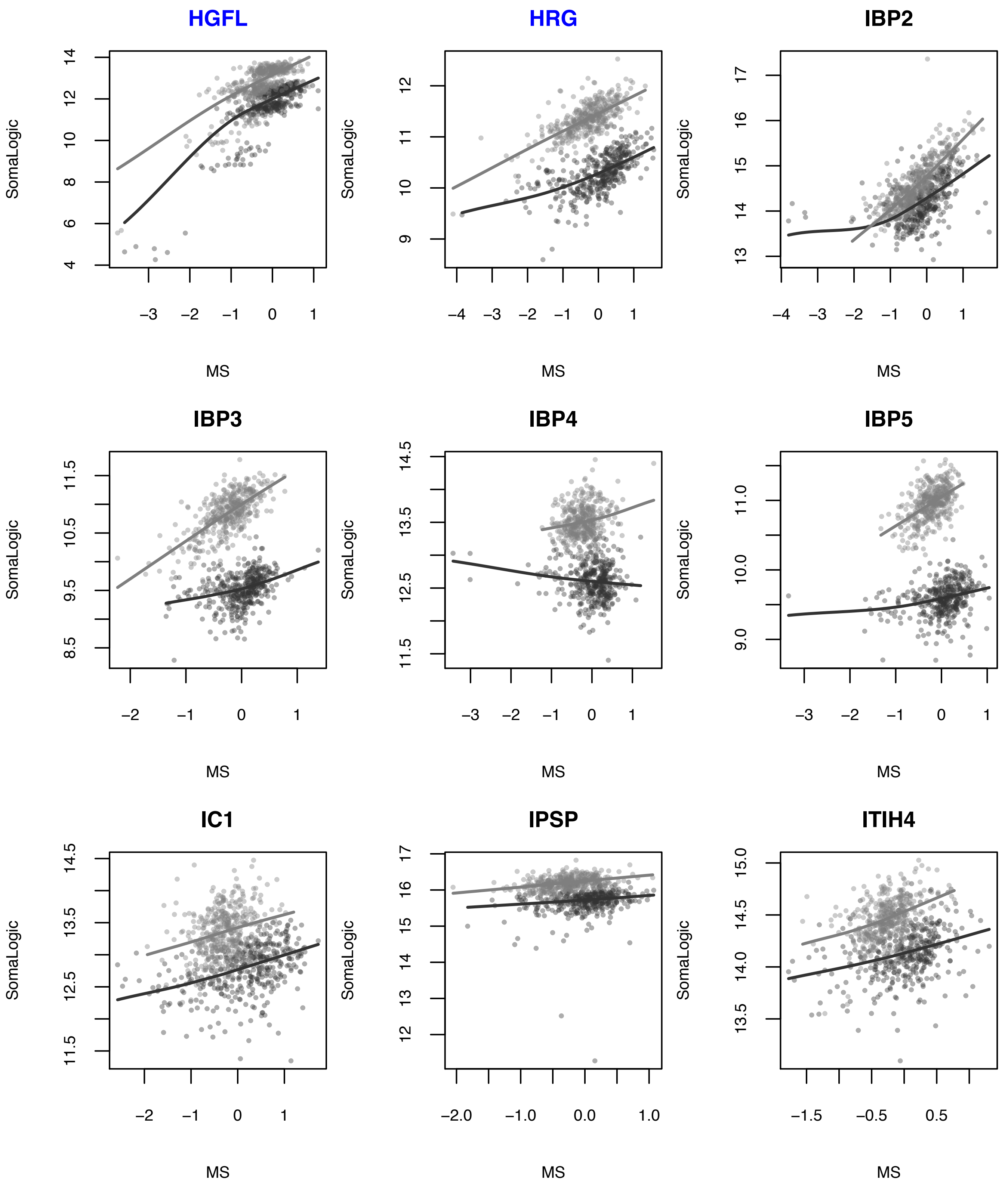
HGFL
SomaLogic
4 6 8 10 12 14
−3 −2 −1 0 1
MS
HRG
SomaLogic
9 10 11 12
−4 −3 −2 −1 0 1
MS
IBP2
SomaLogic
13 14 15 16 17
−4 −3 −2 −1 0 1
MS
IBP3
SomaLogic
8.5 9.5 10.5 11.5
−2 −1 0 1
MS
IBP4
SomaLogic
11.5 12.5 13.5 14.5
−3 −2 −1 0 1
MS
IBP5
SomaLogic
9.0 10.0 11.0
−3 −2 −1 0 1
MS
IC1
SomaLogic
11.5 12.5 13.5 14.5
−2 −1 0 1
MS
IPSP
SomaLogic
12 13 14 15 16 17
−2.0 −1.0 0.0 1.0
MS
ITIH4
SomaLogic
13.5 14.0 14.5 15.0
−1.5 −0.5 0.5
MS

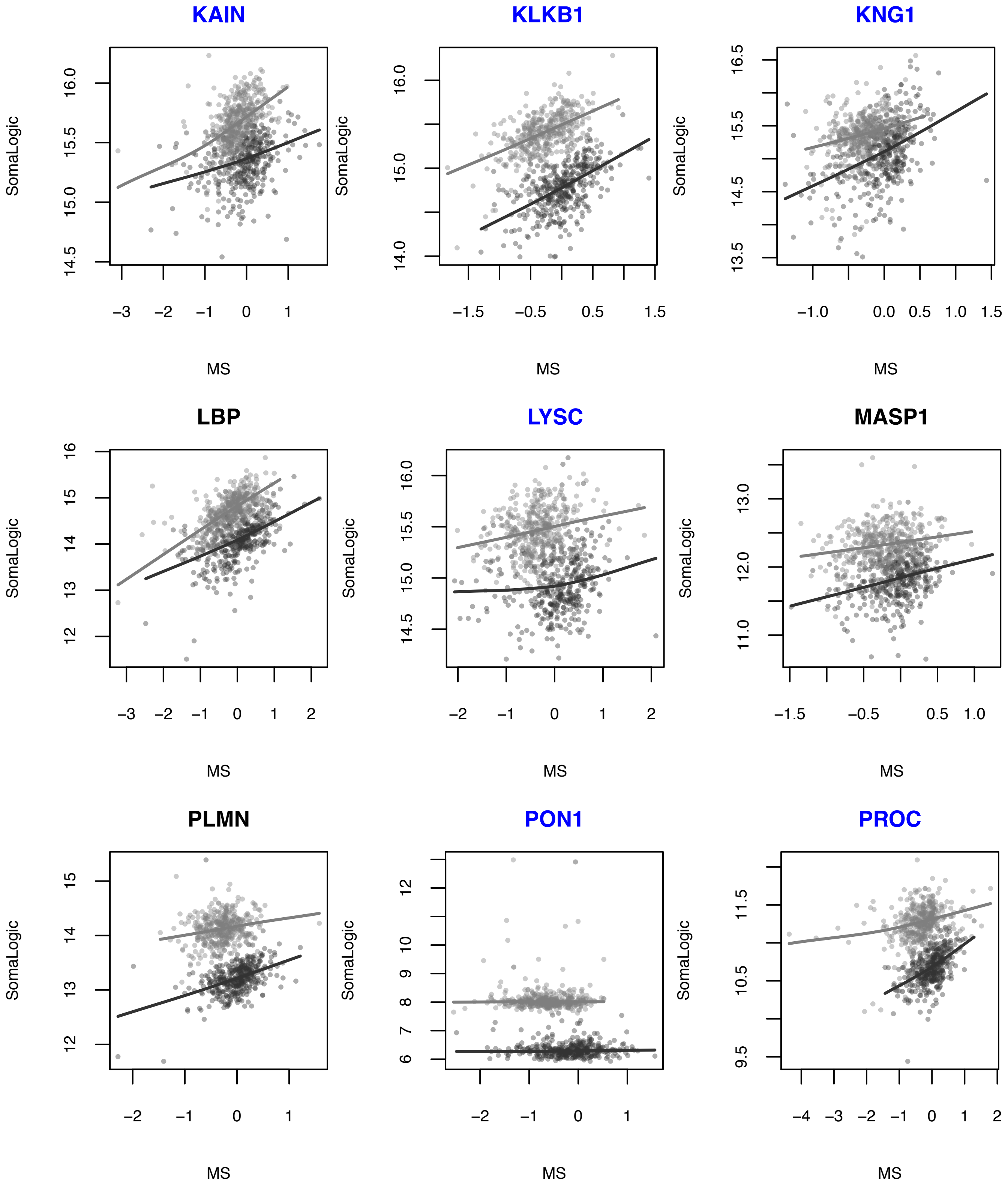
KAIN
KLKB1
KNG1
LBP
LYSC
MASP1
PLMN
PON1
PROC
SomaLogic
MS

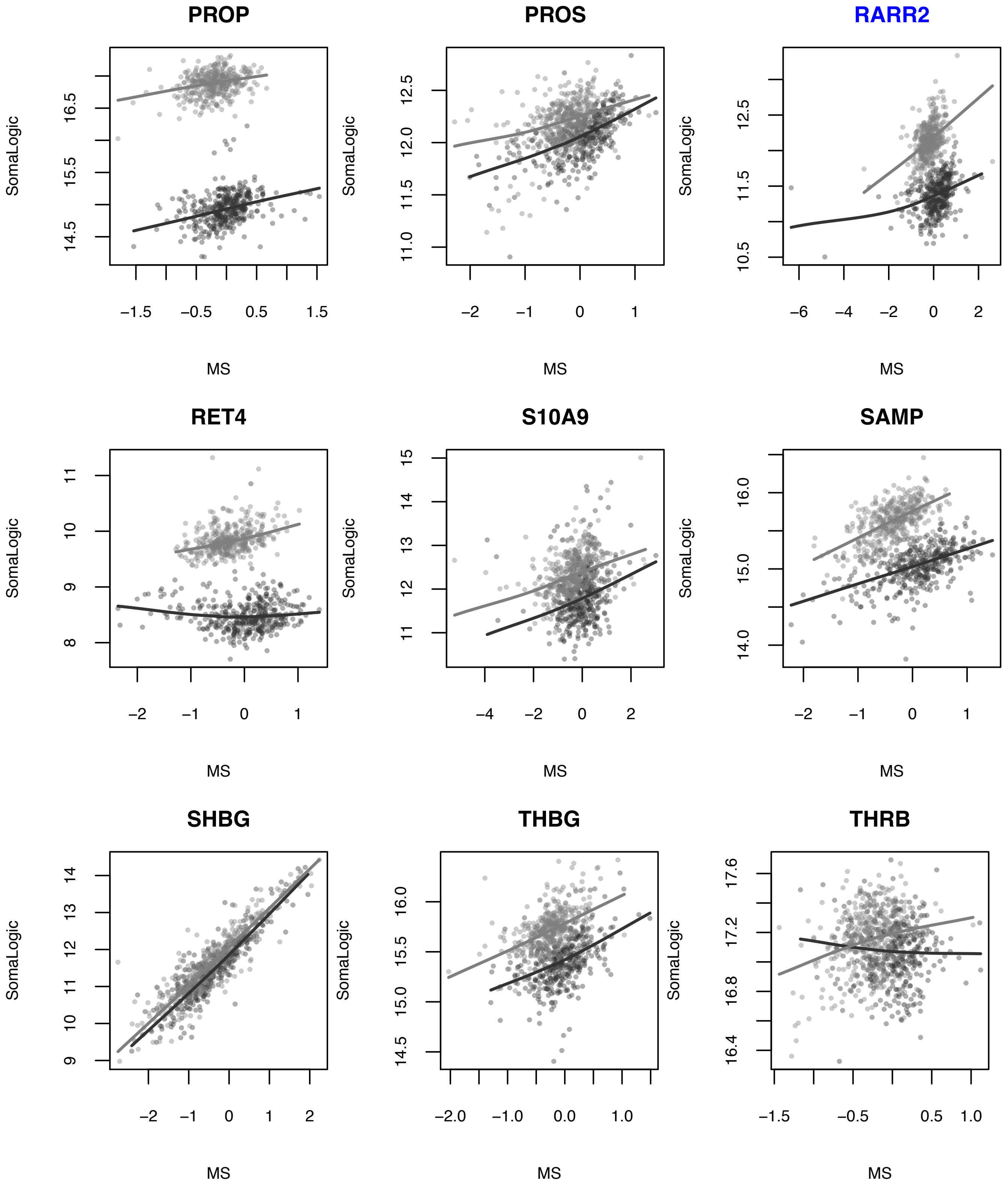
PROP
SomaLogic
14.5 15.5 16.5
−1.5 −0.5 0.5 1.5
MS
PROS
SomaLogic
11.0 11.5 12.0 12.5
−2 −1 0 1
MS
RARR2
SomaLogic
10.5 11.5 12.5
−6 −4 −2 0 2
MS
RET4
SomaLogic
8 9 10 11
−2 −1 0 1
MS
S10A9
SomaLogic
11 12 13 14 15
−4 −2 0 2
MS
SAMP
SomaLogic
14.0 15.0 16.0
−2 −1 0 1
MS
SHBG
SomaLogic
9 10 11 12 13 14
−2 −1 0 1 2
MS
THBG
SomaLogic
14.5 15.0 15.5 16.0
−2.0 −1.0 0.0 1.0
MS
THRB
SomaLogic
16.4 16.8 17.2 17.6
−1.5 −0.5 0.5 1.0
MS
