## Supplementary material for "A fully joint Bayesian quantitative trait locus mapping of human protein abundance in plasma": S2 Appendix

### S2 Appendix to:
### "A fully joint Bayesian quantitative trait locus mapping of human protein abundance in plasma"
### Addendum to the synthetic data generation design and computational performance of LOCUS

#### Emulation of the MS QTL data

Simulation studies can be far from real data scenarios. In an attempt to overcome this, and produce numerical experiments that have may give a realistic account of the performance in real-world studies, we gave a special emphasis to the generation of synthetic data that would best reflect the real data from the discovery study Ottawa.

To this end we used the real SNP data from chromosome one as candidate predictors; the linkage disequilibrium pattern corresponding to the first blocks of SNPs is displayed on Figure 3c. The replication of the MS protein levels requires a particular attention; we used our data generator implemented in our freely available R package `echoseq` to closely reproduce the variance and block correlation structure of the residual protein levels (i.e., the levels before generating associations with the SNPs). Figure A compares the real and simulated protein correlation patterns and indicate a very close agreement between the two.

The effects between the SNPs and the synthetic proteins were also simulated using `echoseq` based on general principles of population genetics (sparsity, distribution of effect sizes, additive dose-effect scheme, natural selection); see Material and methods for details.

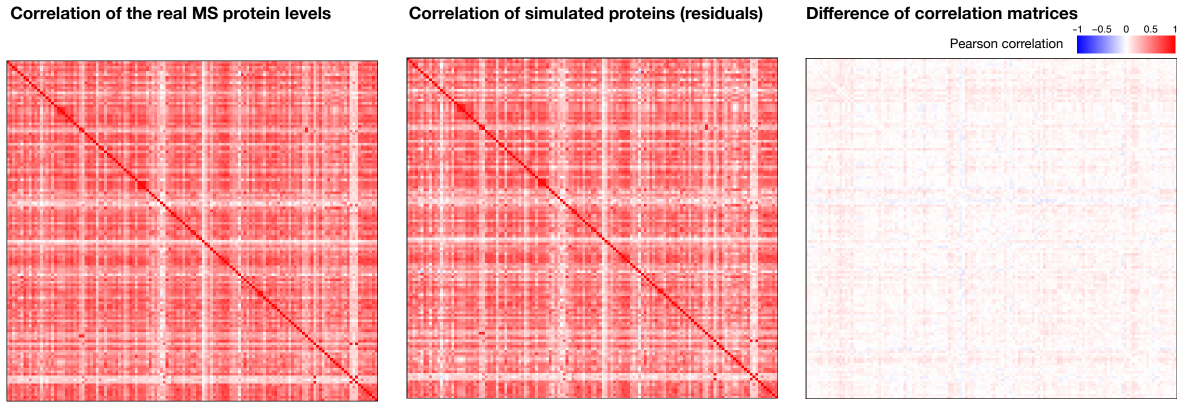


Figure A: **Block correlation structure of the $q = 133$ real and synthetic MS protein levels for the $n = 376$ Ottawa subjects.** Left: Pearson correlation of the MS protein levels; Middle: Pearson correlation of the replicated protein levels; Right: Difference of correlation matrices for the real and simulated protein levels.

#### Computational performance of LOCUS

The runtime of LOCUS for the simulations presented in the paper was similar to that of GEMMA: on average, for one replicate, LOCUS took 5 minutes and 26 seconds to complete, while GEMMA took 7 minutes and 4 seconds, running in parallel on four cores of an Intel Xeon CPU, 2.60 GHz.

Fig B presents runtime profiling for LOCUS when applied to different numbers of SNPs and molecular traits. All runs completed within hours.

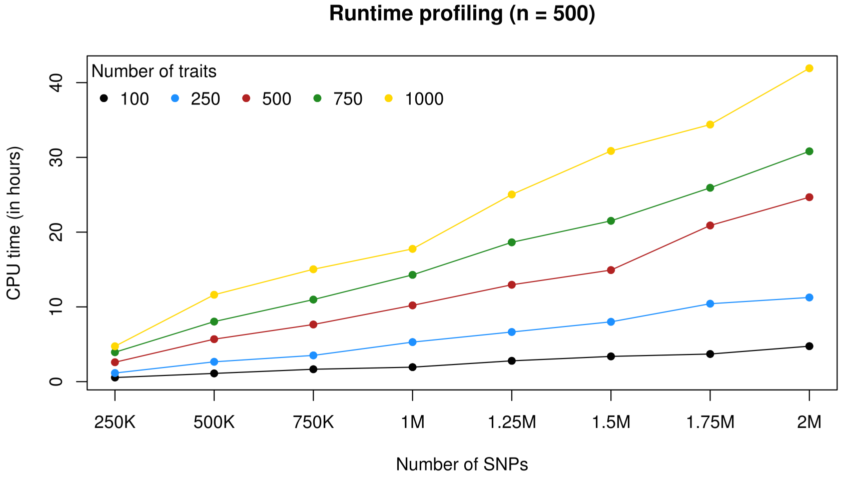


Figure B: **Runtime profiling in CPU hours, for $2.5 \times 10^5$ to $2 \times 10^6$ SNPs and 100 to 1000 traits, on an Intel Xeon CPU at 2.60 GHz with 256 Gb RAM.** For each case, the average runtime of five replications is displayed. The same annealing settings as in the MS and SomaLogic analyses were used (50 geometrically-spaced temperatures and initial temperature $T = 20$).
