## Supplementary material for "A fully joint Bayesian quantitative trait locus mapping of human protein abundance in plasma": S3 Appendix

### S3 Appendix to:
### “A fully joint Bayesian quantitative trait locus mapping of human protein abundance in plasma”
### Addendum to the comparison of LOCUS with the univariate screening analysis GEMMA

#### Six examples of pQTL loci found by LOCUS and missed by the univariate analysis

Fig A displays six examples of hits missed by GEMMA and validated by the LOCUS two-stage MS analysis (loci associated with CO7, ECM1 and ITIH3) and SomaLogic analysis (loci associated with I17RB, PROC and TENA). Each panel shows how some univariate signal is present, of different nature depending on the type of linkage disequilibrium in the loci, but is too weak to be detected after multiplicity correction. In contrast, the multiplicity-adjusted LOCUS analyses could effectively single out and validate individual hits among the strongest GEMMA signals.

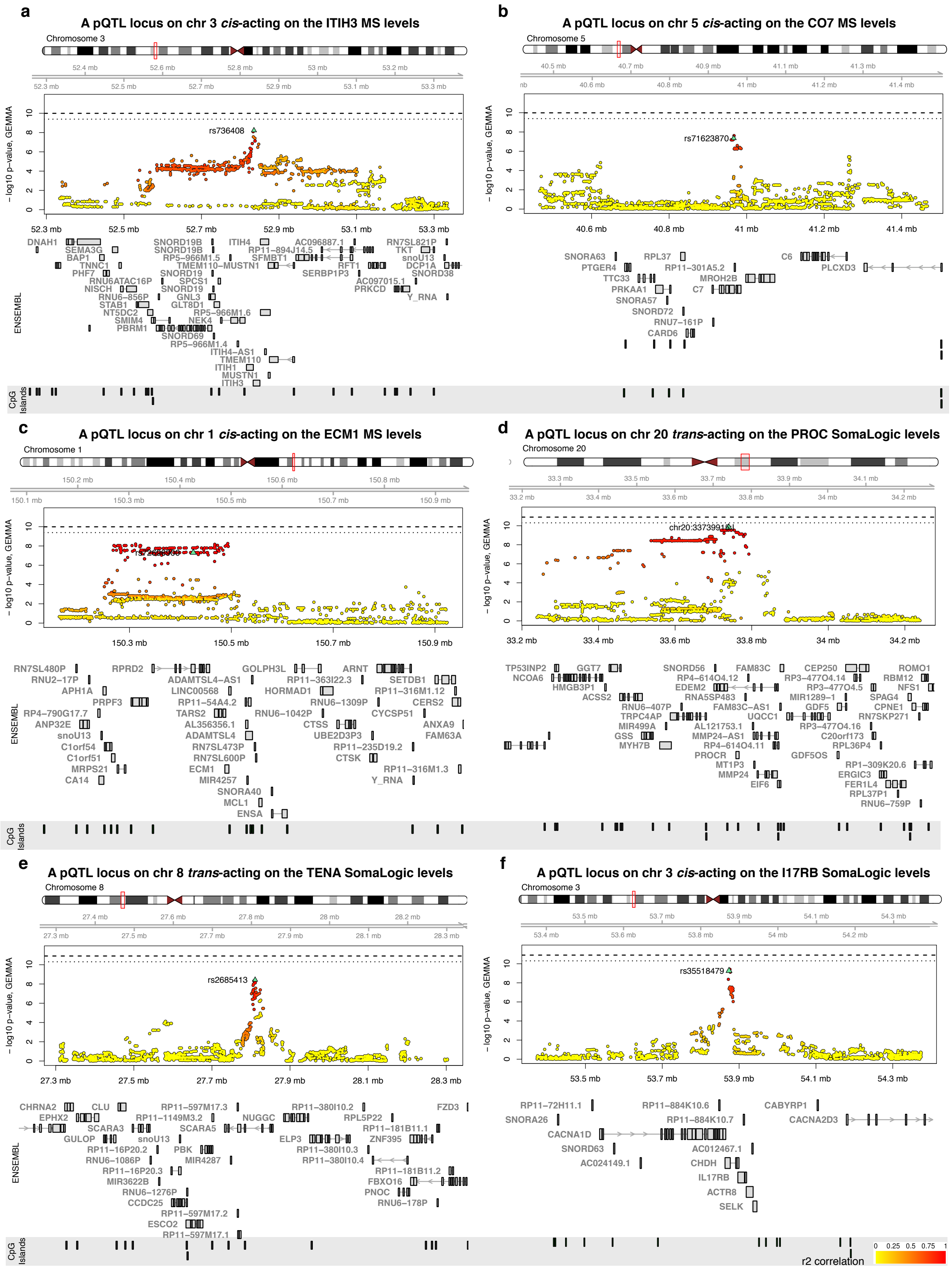


Figure A: **Regional association plots for six loci, identified by the MS (a–c) and SomaLogic (d–f) pQTL analyses.** In each case, the top panel displays the nominal $-\log_{10} p$-values obtained when re-analyzing the Ottawa data with GEMMA [1, 2]; the dashed horizontal line corresponds to a Bonferroni level of $\alpha = 0.05$ and the dotted horizontal line corresponds to a Bonferroni level of $\alpha = 0.2$. The SNP identified by LOCUS is marked with a green triangle, and its correlation in $r^2$ with the surrounding SNPs is indicated by the yellow to red colors. The middle panel shows the transcript positions and the bottom panel shows the CpG island positions.

#### Sensitivity of the univariate analysis to the $p$-value threshold

Fig B displays the replication rates as a function of the $p$-value threshold for individual SNP-protein pairs discovered by GEMMA using the OTTAWA cohort and validated using the DiOGenes cohort, as well as the corresponding numbers of locus-protein pairs for the MS and the SomaLogic analyses. For any choice of threshold, the replication rates are inferior to those of LOCUS two-stage analysis (83% for both the MS and SomaLogic analyses). The number of validated locus-protein pairs is well below that of LOCUS for any reasonable threshold (i.e., $\leq 10^{-6}$) in the MS analysis; this number is similar for GEMMA and LOCUS in the SomaLogic analysis for thresholds $< 10^{-8}$.

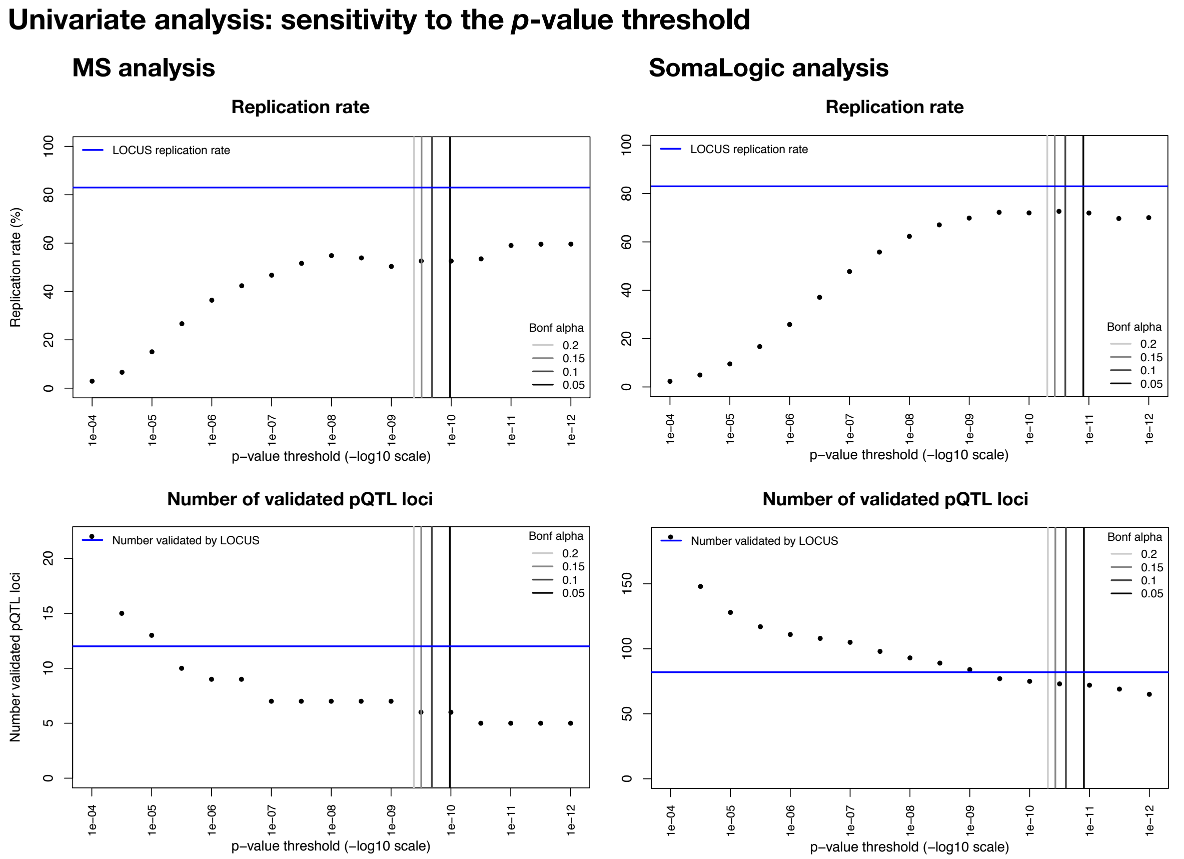


Figure B: **Validation performance of GEMMA univariate analyses as a function of the $p$-value threshold.** Replication rates for pairs SNP-protein (top) and corresponding number of hits summarized by locus-protein (bottom) against $p$-value threshold for the MS (left) and SomaLogic (right) analyses. The performance of LOCUS two-stage analyses is indicated by the blue horizontal line and the discovery Bonferroni thresholds for $\alpha = 0.5, 0.1, 0.15, 0.2$ correspond to the grey vertical lines.
