## Supplementary material for "A fully joint Bayesian quantitative trait locus mapping of human protein abundance in plasma": S4 Appendix

### S4 Appendix to:
### "A fully joint Bayesian quantitative trait locus mapping of human protein abundance in plasma"
### Further examples of pQTL loci with possible implications in metabolic disorders

This appendix provides a general discussion on validated pQTL hits and formulates hypotheses on their clinical relevance. We performed a meta-analysis of the DiOGenes and Ottawa clinical and proteomic data, and found that 35 of the 88 proteins under genetic control had associations with dyslipidemia, insulin resistance or visceral fat-related measurements at FDR 5%; these associations should be attributable metabolic factors independently of overall adiposity, as we controlled for BMI as a potential confounder. They are displayed as a network in Fig A and are listed in Table A. We observed consistent directions of effects in the two cohorts (see Forest plots of Fig 5 and S9 Table for full details). For the proteins from the SomaLogic panel, the above observations may be attributable to the known bias of the SomaLogic assay towards proteins pertaining to inflammation and cancer (S10 Table). However, we also found that the 88 genetically-driven proteins are significantly more associated with the clinical variables compared with protein sets chosen randomly among all proteins quantified by our MS and SomaLogic panels ($p = 0.014$); this enrichment suggests that the primary pQTL analyses can help uncover potential proteomic biomarkers for the Metabolic Syndrome and other obesity-related complications.

| Protein | Protein name | Clin. | SNP | Chr | Position | LOCUS PPI | pQTL valid. $p$-value |
|---|---|---|---|---|---|---|---|
| **CADH5** | **Cadherin-5** | **L** | **rs8176741** | **9** | **136131461** | **1.00** | $\mathbf{4.68 \times 10^{-30}}$ |
| **CD209** | **DC-SIGN** | **L/V** | **rs8176741** | **9** | **136131461** | **1.00** | $\mathbf{7.74 \times 10^{-10}}$ |
| | | | **rs2519093** | **9** | **136141870** | **1.00** | $\mathbf{6.24 \times 10^{-26}}$ |
| CFAB | Factor B | G/L | rs7772063 | 6 | 31906334 | 0.85 | $9.61 \times 10^{-4}$ |
| | | | rs641153 | 6 | 31914180 | 1.00 | $3.92 \times 10^{-12}$ |
| CNTN2 | CNTN2 | L | rs11240396 | 1 | 205205081 | 1.00 | $6.82 \times 10^{-14}$ |
| CO7 | C7 | L | rs71623870 | 5 | 40966676 | 0.83 | $4.03 \times 10^{-4}$ |
| ECM1 | ECM1 | L/V | rs34964511 | 1 | 150298015 | 1.00 | $3.77 \times 10^{-6}$ |
| | | | rs71578487 | 1 | 150340059 | 1.00 | $1.07 \times 10^{-11}$ |
| | | | rs72696900 | 1 | 150425256 | 0.82 | $1.5 \times 10^{-6}$ |
| | | | rs11802612 | 1 | 150427279 | 1.00 | $3.7 \times 10^{-6}$ |
| | | | rs35094010 | 1 | 150449557 | 1.00 | $3.76 \times 10^{-6}$ |
| ESTD | Esterase D | L | rs73193065 | 13 | 47383681 | 0.90 | $2.31 \times 10^{-15}$ |
| FA12 | Coagulation factor XII | L/V | rs55785724 | 5 | 176817583 | 1.00 | $1.34 \times 10^{-5}$ |
| FA7 | Coagulation Factor VII | L/V | rs3093233 | 13 | 113758130 | 1.00 | $3.11 \times 10^{-88}$ |
| FCN2 | FCN2 | L/V | rs3811140 | 9 | 137772111 | 1.00 | $9.66 \times 10^{-14}$ |
| FCN3 | Ficolin-3 | L/V | rs10902652 | 1 | 27558522 | 1.00 | $1.62 \times 10^{-3}$ |
| FETUA | a2-HS-Glycoprotein | G | rs2593813 | 3 | 186332571 | 1.00 | $2.47 \times 10^{-10}$ |
| | | | rs2593813 | 3 | 186332571 | 1.00 | $4.51 \times 10^{-8}$ |
| **HEMO** | **Hemopexin** | **L** | **rs10801560** | **1** | **196714600** | **1.00** | $\mathbf{2.36 \times 10^{-26}}$ |
| I17RA | IL-17 sR | G | rs738035 | 22 | 17594886 | 1.00 | $1.48 \times 10^{-20}$ |
| I17RB | IL-17B R | L/V | rs35518479 | 3 | 53873814 | 0.76 | $9.98 \times 10^{-6}$ |
| IDUA | IDUA | L/V | rs10017289 | 4 | 943534 | 1.00 | $1.22 \times 10^{-11}$ |
| IL18R | IL-18 Ra | L/V | rs3836108 | 2 | 103037742 | 1.00 | $5.22 \times 10^{-26}$ |
| IL1AP | IL-1 R AcP | G/L/V | rs724608 | 3 | 190348810 | 1.00 | $8.7 \times 10^{-114}$ |
| IL6RA | IL-6 sRa | G | rs4845372 | 1 | 154415396 | 1.00 | $1.72 \times 10^{-81}$ |
| ITIH3 | Inter-alpha-trypsin inhibitor heavy chain H3 | L/V | rs736408 | 3 | 52835354 | 0.97 | $1.46 \times 10^{-6}$ |
| KAIN | Kallistatin | L | rs5511 | 14 | 95033595 | 1.00 | $9.9 \times 10^{-24}$ |
| KLKB1 | Prekallikrein | L | rs80177406 | 4 | 187166024 | 0.99 | $3.54 \times 10^{-6}$ |
| KNG1 | Kininogen HMW | L | rs1621816 | 3 | 186439173 | 1.00 | $1.44 \times 10^{-13}$ |
| KYNU | KYNU | G/L/V | rs6741488 | 2 | 143793701 | 1.00 | $3.22 \times 10^{-20}$ |
| **LYAM2** | **sE-Selectin** | **L/V** | **rs2519093** | **9** | **136141870** | **1.00** | $\mathbf{6.81 \times 10^{-62}}$ |
| LYSC | Lysozyme | L | rs71094714 | 12 | 69790495 | 1.00 | $8.41 \times 10^{-19}$ |
| MPRI | IGF-II receptor | L | rs3777411 | 6 | 160476945 | 1.00 | $4.95 \times 10^{-11}$ |
| PA2GA | NPS-PLA2 | G/L/V | rs6672057 | 1 | 20293791 | 1.00 | $3.86 \times 10^{-15}$ |
| PCSK7 | PCSK7 | L/V | rs11216284 | 11 | 117003060 | 1.00 | $8.17 \times 10^{-31}$ |
| **PROC** | **Protein C** | **L/V** | **rs141091409** | **20** | **33739915** | **0.43** | $\mathbf{1.66 \times 10^{-18}}$ |
| RARR2 | TIG2 | G/L/V | rs1047586 | 7 | 150035459 | 0.96 | $2.39 \times 10^{-11}$ |
| SIGL6 | Siglec-6 | L | rs8101887 | 19 | 52029477 | 1.00 | $3.39 \times 10^{-14}$ |
| SPRL1 | SPARCL1 | L/V | rs7681694 | 4 | 88462729 | 0.99 | $5.70 \times 10^{-14}$ |
| **TXD12** | **TXD12** | **L** | **rs13062429** | **3** | **49559485** | **1.00** | $\mathbf{2.26 \times 10^{-5}}$ |
| | | | **rs34519883** | **3** | **49575831** | **1.00** | $\mathbf{5.39 \times 10^{-33}}$ |
| WFKN2 | WFKN2 | G/L/V | rs9303566 | 17 | 48922281 | 1.00 | $3.38 \times 10^{-11}$ |

Table A: **Proteins associated with clinical parameters (Fig A) and controlled by pQTL variants.** All associations were detected at FDR $< 5\%$. Associations with glycemic traits (fasting glucose, insulin, HOMA-IR) are indicated by $G$, with total lipid traits (HDL, LDL, triglycerides, total cholesterol), by $L$, and with visceral fat (visceral adiposity index), by $V$. *Trans*-pQTL associations are in bold.

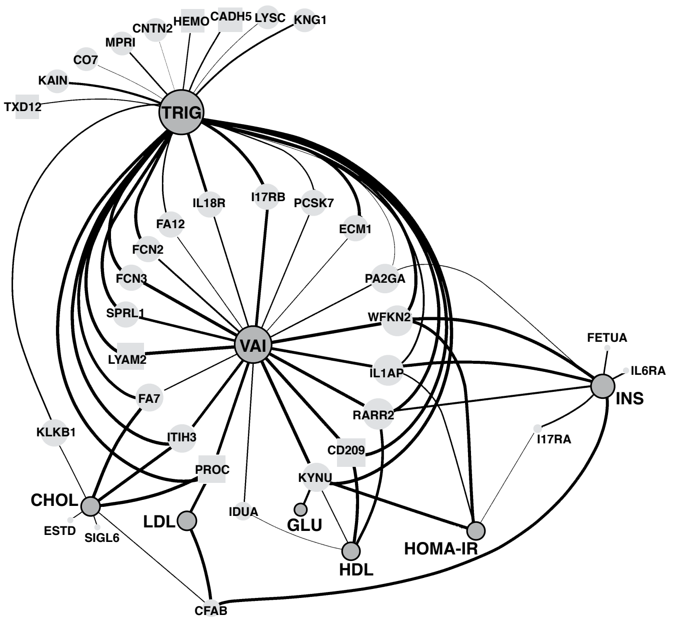


Figure A: **Associations of proteins under genetic control with clinical parameters.** Network displaying the associations (FDR < 5%) between protein levels and clinical variables obtained by meta-analysis, adjusting for age, gender and BMI. The nodes representing clinical parameters are in dark grey with black borders (fasting glucose, HDL, HOMA-IR, insulin resistance, LDL, total cholesterol, triglycerides, visceral adiposity index), and those representing proteins are in light grey with the type of genetic control, *cis* or *trans*, depicted with circles or squares, respectively. The edge thickness is proportional to the significance of association, and the node size is proportional to its connectivity.

As shown in the network of Fig A, the triglyceride measurements and visceral adiposity index (VAI) have the highest degree of connectivity and are connected with measures of insulin resistance and other lipid traits via proteins such as FA7, IL1AP, KYNU, PROC, RARR2 and WFKN2. CFAB, FETUA, PA2GA have lower connectivity, yet are relevant in the context of obesity [1–3]. *Trans*-regulated proteins (Fig A) were also implicated in clinical associations: CADH5, CD209 and LYAM2, all controlled by the pleiotropic locus *ABO*; HEMO (Hemopexin), a liver glycoprotein controlled by the *CFH* locus, itself coding for another liver glycoprotein; PROC controlled by its own receptor *PROCR*; and TXD12 (thioredoxin domain containing 12), controlled by the *DAG1/BSN* locus.

In the subsequent sections, we expand on the possible functional and biomedical relevance of several representative examples of pQTL associations in the context of obesity complications. Unless otherwise specified, all associations described have meta-analysis FDR corrected $p$-value below 5%, and we provide their nominal $p$-values in parentheses.

#### Clinical associations with proteins controlled by the pleiotropic locus *ABO*

Our clinical analyses indicated associations of CD209 and LYAM2 with triglycerides and visceral fat, and CADH5 with triglycerides only (Fig 5). The LYAM2 levels were associated with all the glycemic variables in the Ottawa cohort (fasting glucose: $p = 6.43 \times 10^{-6}$, fasting insulin: $p = 3.54 \times 10^{-4}$,

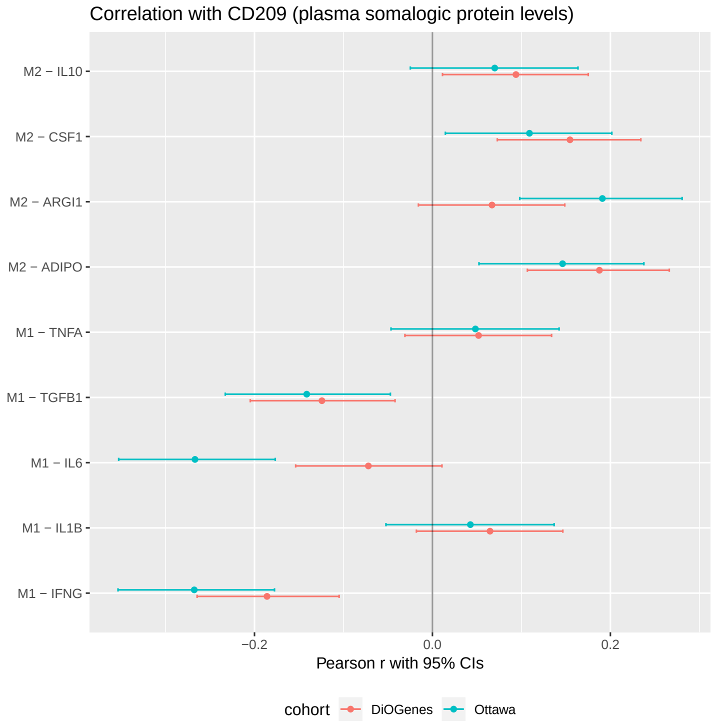


Figure B: **Correlation between CD209 and other macrophage protein levels.** Pearson correlation coefficient with 95% confidence interval for the association between CD209 protein levels (controlled by $rs8176741$, see Results section) and M1-M2 macrophage protein levels in Ottawa and in DiOGenes.

HOMA-IR: $p = 1.8 \times 10^{-4}$), but only with fasting glucose in the DiOGenes cohort ($p = 8.91 \times 10^{-4}$), although we observed a trend for HOMA-IR (nominal $p = 0.02$, corrected $p = 0.15$). Since the Ottawa subjects are more insulin-resistant than the DiOGenes subjects (average HOMA-IR with standard deviation: 4.97(3.88) versus 3.00(1.71), $p = 2.52 \times 10^{-18}$; S1 Table), LYAM2 might represent a marker of insulin-resistance severity. Consistent with this hypothesis, the plasma levels of LYAM2 are employed as a biomarkers of endothelial dysfunction and risk of type 2 diabetes [4].

We found that CD209 circulating levels were positively associated with HDL, negatively with triglyceride levels, and, consistently with these effects, negatively with visceral fat index (Fig 5), suggesting beneficial effects of high CD209 levels. Further investigation using deconvolution of adipose tissue gene expression profiles showed that CD209 is predominantly secreted by M2 macrophages [5]. These cells are involved in extracellular matrix remodelling and secrete cytokines with an anti-inflammatory role, counteracting the effect from pro-inflammatory macrophages M1. Interestingly, other adipose cell types, including M1 macrophages display little, if any, CD209 expression. M1 and M2 macrophages have been extensively discussed in the context of obesity, and it is well established that M2 macrophages have a protective role against obesity and insulin resistance [6] and increase fatty oxidation and oxidative phosphorylation [7]. In our data (both the Diogenes and Ottawa cohorts, and at FDR 5%), we found that CD209 levels were positively correlated with M2 secreted proteins (IL10, CSF1, ARGI1) and negatively correlated with M1 pro-inflammatory markers, such as TGF-beta, IL6 and interferon-gamma (Fig B).

Importantly, CD209 was positively associated with circulating levels of adiponectin, an hormone secreted in adipose tissue which plays a key role in glucose regulation, fatty acid oxidation and triglycerides clearance. This lends support that CD209 could be a secreted protein, released by M2 macrophages from adipose tissue, with a beneficial role in controlling lipid levels, thereby possibly protecting from developing dyslipidemia and related metabolic complications.

#### Clinical associations with the XRCC6 protein levels

We observed significant associations between the XRCC6 protein levels and several clinical variables in the Ottawa cohort (FDR $< 5\%$). Higher expression was associated with decreased HDL ($p = 5.83 \times 10^{-4}$), as well as with higher triglycerides ($p = 4.39 \times 10^{-4}$), insulin levels ($p = 4.50 \times 10^{-4}$) and visceral adiposity ($p = 5.94 \times 10^{-5}$; Fig 5). We only found marginal associations using the DiOGenes data for insulin levels (nominal $p = 0.02$, corrected $p = 0.14$) and HOMA-IR (nominal $p = 0.02$, corrected $p = 0.16$). The directionality of these effects was consistent in both cohorts. As the Ottawa subjects were more severely obese, the effects might be larger for subjects with pronounced Metabolic Syndrome.

Our pQTL sentinel SNP, rs4756623, is intronic and located within the *LRRC4C* gene, a binding partner for Netrin G1 and member of the axon guidance [8]. To our knowledge, *LRRC4C* has not been previously described in the context of obesity, insulin resistance or type 2 diabetes. However, its partner Netrin G1 is known to promote adipose tissue macrophage retention, inflammation and insulin resistance in obese mice [9]. The underlying regulatory mechanisms between rs4756623 and the *XRCC6* locus should be clarified, and functional studies will be required to understand their physiological impact.

#### CFAB and RARR2, mediators of adipogenesis are under genetic control

CFAB (complement factor B) and RARR2 (Retinoic acid receptor responder protein 2) levels associate with distinct clinical parameters (Figs A and 5), yet both play a role in adipogenesis and hence are particularly interesting in the context of obesity and related co-morbidities.

The CFAB protein controls the maturation of adipocytes in rat [1] and has a determinant role in metabolic and cardiovascular dysfunctions linked with the Metabolic Syndrome [10]. In our study, both the MS and SomaLogic measurements were positively associated with BMI (MS: $p = 2.08 \times 10^{-8}$, SomaLogic: $p = 2.23 \times 10^{-13}$) and with fasting insulin (adjusting for BMI; MS: $p = 4.45 \times 10^{-5}$, SomaLogic: $p = 3.44 \times 10^{-4}$). The CFAB SomaLogic levels were negatively associated with cholesterol ($p = 1.43 \times 10^{-3}$), LDL ($p = 1.30 \times 10^{-5}$), and with HDL at higher FDR (nominal $p = 1.47 \times 10^{-2}$, corrected $p = 0.11$). This is consistent with previous work linking *CFB* gene expression from different human adipose tissue fractions with insulin resistance and lipid levels [11].

Our MS and SomaLogic analyses independently highlighted the same *cis*-acting locus as putative regulator of the CFAB protein. In particular, the sentinel pQTL SNP detected in the SomaLogic analysis, rs641153, is a missense variant located in the MHC region, 180 base pairs away from a transcription binding site (significantly closer than other SNPs, $p = 1.16 \times 10^{-2}$). Further investigation using JASPAR [12] and SNP2TFBS [13]indicated that rs641153 may affect the binding sites of four transcription factors (EBF1, TFAP2A, TFAP2C and HNFA), and this SNP has indeed been reported as both e- and pQTL [14, 15].

RARR2 (Chemerin protein) is encoded by an essential adipogenesis gene, *RARRES2*. This adipokine plays an important role in inflammation, adipogenesis, angiogenesis and glucose homeostasis [16, 17]. Our pQTL analyses indicated a *cis* association between a missense variant, rs1047586, and RARR2 protein levels, in line with previous findings reporting this SNP as e- and pQTL (S4 and S7 Tables). Moreover, our analyses of protein levels revealed significant associations with triglycerides, fasting insulin and HDL (Fig 5, S9 Table), which is consistent with previously described pleiotropic associations of *RARRES2* variants with circulating RARR2, triglyceride levels and diverse measurements related to inflammation [18], and findings from animal models [19, 20]. Moreover, MS and SomaLogic RARR2 levels were strongly associated with visceral fat, even when controlling for BMI (Fig 5; S9 Table), further strengthening the relevance of this protein in the development of the Metabolic Syndrome [17, 21].

#### The importance of IL1AP for Metabolic Syndrome

The IL-1 pathway plays a critical role in the immune-response associated with obesity and type 2 diabetes [22]; other IL-1 related cytokines, such as IL-1ra, are also well documented in the context of type 1 and type 2 diabetes [23]. The IL1AP (IL-1 receptor accessory) protein is a co-receptor of the IL-1 receptor, and its soluble levels were found reduced in obese subjects [24]. Our analyses found an association between rs724608 and IL1AP, corroborating previously identified associations with SNPs in LD ($r^2 = 0.93$) [24].

We found associations between IL1AP expression and measures of fasting insulin levels ($p = 3.88 \times 10^{-5}$), HOMA-IR ($p = 3.89 \times 10^{-4}$), triglycerides ($p = 1.61 \times 10^{-3}$) and visceral fat ($p = 2.1 \times 10^{-4}$) (Figs A and 5). Moreover, worsened Metabolic Syndrome score [25] were associated with lower protein levels ($p = 1.20 \times 10^{-3}$ in Ottawa and $p = 2.50 \times 10^{-4}$ in DiOGenes).

#### WFKN2, a TGF$\beta$-activity protein with protective effect against metabolic disorders

The role of the WFKN2 protein and of its coding gene, *WFIKKN2*, in regulating TGF$\beta$ activity has been extensively studied in muscle and skeletal muscle [26], but, to our knowledge, not in other tissues. We describe it for the first time in the context of obesity and metabolic disorders. We found that higher protein levels were associated with lower levels of fasting insulin, triglycerides, HOMA-IR and visceral fat (Fig 5), suggesting a protective role against metabolic dysregulation.

Our analyses suggested that the WFKN2 levels are controlled by rs9303566, which is consistent with other p- and eQTL studies (S4–S7 Tables). This SNP was found to be associated with DNA methylation and histone marks [27, 28], and is located within 100 base pairs of a transcription factor binding site, with numerous factors such as MYBL2, NFIC, EP300 and MXI1. It is in strong LD with other SNPs with potential regulatory impact; for instance, it is located 9Kb upstream to rs8072476 ($r^2 = 0.97$), which overlaps another cluster of transcription factor binding sites (FOXA1, ESR1, USF1 & 2, TFAP2A & 2C).

#### Inflammation mediated proteins and their role in insulin resistance

We found a *cis* effect of rs6741488 on KYNU (Kynureninase) plasmatic levels. KYNU is an enzyme involved in the biosynthesis of nicotinamide adenine dinucleotide (NAD) cofactors from tryptophan. This protein and its pathway have been found to be particularly relevant for obesity and associated metabolic disorders. KYNU was found to be up-regulated by pro-inflammatory cytokines in human primary adipocytes, and more so in the omental adipose tissue of obese compared to lean control subjects [29]. Other studies indicated that the kynurenine pathway (KP) may act as an inflammatory sensor, and that increased levels of its catabolites may be linked with several cardiometabolic defects, including cardiovascular disease, diabetes and obesity [30]. In our cohorts, higher KYNU levels were associated with decreased HDL levels ($p = 6.66 \times 10^{-4}$), and increased triglycerides levels ($p = 3.43 \times 10^{-8}$), visceral fat ($p = 2.51 \times 10^{-8}$) and insulin resistance (marginally, nominal $p = 2.53 \times 10^{-2}$, corrected $p = 0.17$), see Fig 5; as expected, higher protein levels were associated with a worsened Metabolic Syndrome score (Ottawa $p = 8.23 \times 10^{-5}$; DiOGenes $p = 3.62 \times 10^{-6}$).

Recent work suggested a causal link between obesity and cancer, mediated by KP activation through inflammatory mechanisms [31]. Interestingly, our analyses highlighted two soluble interleukin receptor antagonist proteins, namely IL6RA and I17RA, that were both under genetic control and associated with insulin resistance (Fig A). We did not find significant correlation between the I17RA and KYNU protein levels, but we did observe a significant negative correlation between IL6RA and KYNU (Ottawa $p = 0.01$ and DiOGenes $p = 4 \times 10^{-3}$). We found a link between the plasma levels of KYNU and pro-inflammatory molecules, namely, IL6, IFNG and TNF$\alpha$. In the Ottawa cohort, where subjects displayed high low-grade inflammation status, KYNU was positively associated with IL6 and IFNG at FDR 5%, while in DiOGenes, we found a positive association with IFNG only (see Fig C). Finally, metabolic dysfunctions mediated via KP may relate to another inflammatory pathology, namely, psoriasis [32], a skin disease aggravated by obesity and improved by weight loss [33, 34].

Our results thus highlighted pQTLs with probable roles in inflammation and subsequent metabolic dysfunctions, reinforcing previous discussion [30, 35] of the potential of KP therapeutic inhibitors against cardiovascular disease and metabolic disorders.

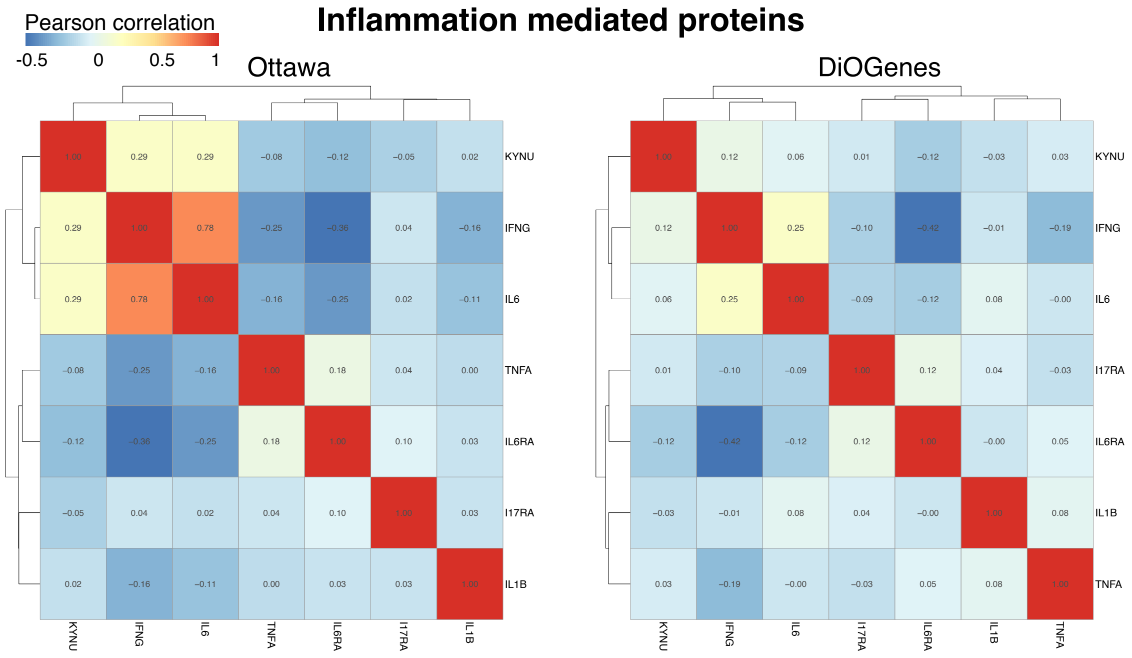


Figure C: **Correlation of KYNU, IFNG, IL6, THFA, IL6RA, I17RA and IL1B in Ottawa (left) and in DiOGenes (right).**

#### Complement/coagulation: a *trans*-acting insertion linking PROC and its receptor

PROC (Protein C, coding gene PROC on chromosome 2) and its paralog protein FA7 (Coagulation Factor 7, coding gene F7 on chromosome 13) regulate the complement and the coagulation systems. Both systems promote inflammation [36] and contribute to metabolic dysfunction in the adipose tissue and liver [37]. Our analyses suggested novel pQTLs for these proteins (S3 Table): FA7 was associated with rs3093233, which is a known eQTL of *F7* and *F10* in several tissues (S7 Table). PROC may be controlled by *trans*-regulatory mechanisms, initiated in its receptor gene, *PROCR*, on chromosome 20; it was indeed associated with an insertion, rs141091409, located 20Kb upstream of *PROCR*, an association observed with both our proteomic platforms. Previous studies found associations between cardiovascular disease and variants located in the *PROC* or *PROCR* genes [38, 39]. Interestingly, our hit, rs141091409, was in strong LD ($r^2 > 0.95$) with the missense variant rs867186, previously identified as associated with coronary heart disease [39].

Our clinical analyses support the relation of PROC and FA7 levels with lipid traits: both were positively associated with cholesterol, triglycerides and visceral fat (Figs A and 5). PROC levels were quantified by both platforms, and displayed consistent results. The SomaLogic measurements of PROC were positively associated with LDL ($p = 5.39 \times 10^{-5}$). The role of these proteins for cardiovascular disease and NAFLD diseases in the overweight/obese population would merit further investigation.

[11] Moreno-Navarrete, J. M., Martínez-Barricarte, R., Catalán, V., Sabater, M., Gómez-Ambrosi, J., Ortega, F. J., Ricart, W., Blüher, M., Frühbeck, G., Rodríguez de Cordoba, S., et al. Complement

factor H is expressed in adipose tissue in association with insulin resistance. *Diabetes*, 59:200–209, 2010.

[18] Er, L., Wu, S., Hsu, L., Teng, M., Sun, Y., and Ko, Y. Pleiotropic Associations of RARRES2 Gene Variants and Circulating Chemerin Levels: Potential Roles of Chemerin Involved in the Metabolic and Inflammation-Related Diseases. *Mediators of Inflammation*, 2018, 2018.

[34] Jensen, P., Christensen, R., Zachariae, C., Geiker, N. R. W., Schaadt, B. K., Stender, S., Hansen, P. R., Astrup, A., and Skov, L. Long-term effects of weight reduction on the severity of psoriasis

in a cohort derived from a randomized trial: a prospective observational follow-up study. *The American Journal of Clinical Nutrition*, 104:259–265, 2016.
