## Supplementary material for "A fully joint Bayesian quantitative trait locus mapping of human protein abundance in plasma": S5 Appendix

### S5 Appendix to:
### "A fully joint Bayesian quantitative trait locus mapping of human protein abundance in plasma"
#### Addendum to the genotyping quality control

The same quality control steps have been applied to the Ottawa and DiOGenes genotyping datasets. In the Ottawa study, $2,418$ subjects had genotyping data available and 80 were discarded for quality reasons (3.3%). In the DiOGenes study, 833 subjects had genotyping data available and 25 (3%). The numbers of subjects removed by the filters are detailed in Table A. These numbers concern subjects with genotyping data; only a subset of them have proteomic data available and could be used for our pQTL analyses, see Material and methods.

| Filter | Ottawa study | DiOGenes study |
|---|---|---|
| Low call rate ($< 95\%$) | 7 | 2 |
| Abnormal autosomal heterozygosity | 28 | 11 |
| XXY karyotype | 2 | 1 |
| Gender discrepancies | 41 | 9 |
| High IBS | 11 | 6 |

Table A: Number of subjects discarded by the different quality filters for the Ottawa and DiOGenes studies. A same subject can be flagged by several filters.
